# Actively replicating gut bacteria identified by 5-ethynyl-2’-deoxyuridine (EdU) click chemistry and cell sorting

**DOI:** 10.1101/2022.07.20.500840

**Authors:** Eve T. Beauchemin, Claire Hunter, Corinne F. Maurice

**Author notes:** Corresponding Author: Corinne F. Maurice.

## Abstract

The composition of the intestinal bacterial community is well described, but recent research suggests that the metabolism of these bacteria plays a larger role in health than which species are present. One fundamental aspect of gut bacterial metabolism that remains understudied is bacterial replication. Indeed, there exist few techniques which can identify actively replicating gut bacteria. In this study, we aimed to address this gap by adapting 5-ethynyl-2’-deoxyuridine (EdU) click chemistry (EdU-click), a metabolic labeling method, coupled with fluorescence-activated cell sorting and sequencing (FACS-Seq) to characterize replicating gut bacteria. We first used EdU-click with human gut bacterial isolates and show that many of them are amenable to this technique. We then optimized EdU-click and FACS-Seq for murine fecal bacteria and reveal that *Prevotella* UCG-001 and *Ileibacterium* are enriched in the replicating fraction. Finally, we labelled the actively replicating murine gut bacteria during exposure to cell wall-specific antibiotics *in vitro*. We show that regardless of the antibiotic used, the actively replicating bacteria largely consist of *Ileibacterium*, suggesting the resistance of this taxon to perturbations. Overall, we demonstrate how combining EdU-click and FACSeq can identify the actively replicating gut bacteria and their link with the composition of the whole community in both homeostatic and perturbed conditions. This technique will be instrumental in elucidating *in situ* bacterial replication dynamics in a variety of other ecological states, including colonization and species invasion, as well as for investigating the relationship between the replication and abundance of bacteria in complex communities.

## Introduction

The community of microorganisms which reside in the intestines of animals, called the gut microbiota, forms one of the densest and most diverse microbial assemblages in the world. The gut microbiota primarily consists of bacteria and the viruses that infect them, with lesser numbers of archaea, fungi, and other eukaryotic microorganisms.^1–4^ Due to their numerical dominance and the existence of tools to study them, bacteria are the best characterized microorganisms in the gut. Among many other roles, these bacteria help to train the immune system, prevent pathogen invasion, and break down indigestible fibers into host-accessible metabolites^4–7^.

Most studies on the gut microbiota to date have focused on describing which bacteria are present in the gut and tried to decipher their links with disease. In recent years, however, more studies have been focusing on additionally describing the activity of these microorganisms.^1, 8–12^ Indeed, the metabolism of the gut bacteria has been suggested to play a bigger role in determining the health status of their host than composition alone.^1, 13^ Furthermore, studying the *in situ* physiology of bacterial communities is suggested to provide a more detailed and accurate picture of how microbiotas function and interact within themselves and with their (host) environment, as compared to culture- or sequence-based methods.^14^

One aspect of microbial physiology which is of growing interest is bacterial replication. The replication strategies of individual gut microbial members have been shown to impact the overall functional capacity of the community and could allow the description and prediction of the outcomes of ecological processes such as competition, succession, and extinction events.^15, 16^ Several techniques in the fields of systems biology and bioinformatics have recently been developed to quantify the replication dynamics of specific gut bacterial taxa or all bacterial members within the gut community, respectively.^17–19^ Notably, the bioinformatics-based studies report correlations between bacterial replication rates and diverse host health parameters, including the occurrence of type II diabetes and inflammatory bowel disease^20, 21^. Furthermore, some of the systems biology approaches have shown that *Escherichia coli* replication dynamics in the gut are altered during host inflammation,^19^ and that a murine *E. coli* strain successfully colonizes the gut in part by dividing twice as fast as it is removed.^18^

However, there exist fewer physiological methods for measuring the replication of individual bacteria in complex communities *in situ*. The recently developed fluorescent labeling of newly synthesized bacterial peptidoglycan is a purely qualitative approach, which also requires the creation of modified probes to allow for fluorescent *in situ* hybridization to taxonomically identify labeled bacteria, a non-trivial task.^22–25^ Quantitative stable isotope probing of bacterial DNA (DNA-qSIP) is time- and resource-intensive.^16^ However, it is compatible with downstream methods to separate and sequence the DNA of labeled cells, which allows the replication rates of individual taxa to be estimated.^16^ This process of labeling, sorting, and characterizing active bacteria is becoming more and more popular due to its unparalleled ability to link bacterial metabolism and identity.^14, 26^

One physiological method for labeling replicating bacteria, which has not yet been used with the gut microbiota, are thymidine analogues. New thymidine analogues which use fluorescent probes are easier and safer to use than their older radioactive counterparts, especially in the context of probing environmental communities *in situ*.^27, 28^ This is especially the case for the thymidine analogue 5-ethynyl-2’-deoxyuridine (EdU), which uses the reliable, quick, and simple method of ‘click’ chemistry (EdU-click) to fluorescently label incorporated EdU in newly created strands of DNA .^27^ Like qSIP, it can be paired with downstream methods to sort and sequence the labeled cells, providing taxonomic information on the replicating and non-replicating communities. To date, however, EdU-click has only been used once in a complex aquatic microbial community. In this context, the researchers identified that between 4.4 to 49% of bacteria in coastal waters were replicating, with a low number of cells contributing to the majority of the fluorescence signal.^29^ If modified, this approach could be used to investigate the replication dynamics of the gut microbiota. For instance, this technique could help determine whether bacterial replication is a phylogenetic characteristic in the gut, as it is suggested to be in soil communities,^30^ or whether it is a dynamic feature of bacterial metabolism which can change with fluctuating environmental conditions.

We here describe the use of the EdU-click technique first in select gut bacterial isolates, followed by optimization of the procedure for the murine gut microbiota. We then combine this technique with fluorescence-activated cell sorting (FACS) and 16S amplicon sequencing (FACS-Seq) to physically separate and taxonomically identify the replicating bacterial communities in healthy mice. We end by demonstrating one potential application of the optimized EdU-click and FACS-Seq procedure, to identify the replicating gut bacteria after an *in vitro* antibiotic exposure. Our findings reveal that many gut bacterial isolates are amenable to EdU-click, and that the replicating gut bacteria in healthy mice is dominated by *Ileibacterium*. We additionally report that exposure of mouse fecal bacteria to an antibiotic either targeting Gram-negative or Gram-positive bacteria results in an increased presence of *Ileibacterium* in the replicating fraction of the gut.

## Results

### Optimization of EdU-click in gut bacterial isolates

To determine whether EdU could be incorporated into phylogenetically distinct bacteria, we first performed EdU-click followed by flow cytometry (**Fig. 1a**) on nine gut bacterial isolates, spanning five phyla and the two cell wall types: *Bacteroides caccae, Bacteroides fragilis*, and *Prevotella intermedia* (Bacteroidetes, Gram -)*; Clostridium beijerinckii*, *Enterococcus faecalis*, and *Ruminococcus bromii* (Firmicutes, Gram +); *Bifidobacterium longum* subsp. *infantis* (Actinobacteria, Gram +); *Escherichia coli* K12 (Proteobacteria, Gram -); and *Akkermansia muciniphila* (Verrucomicrobia, Gram -).

**Figure 1.**
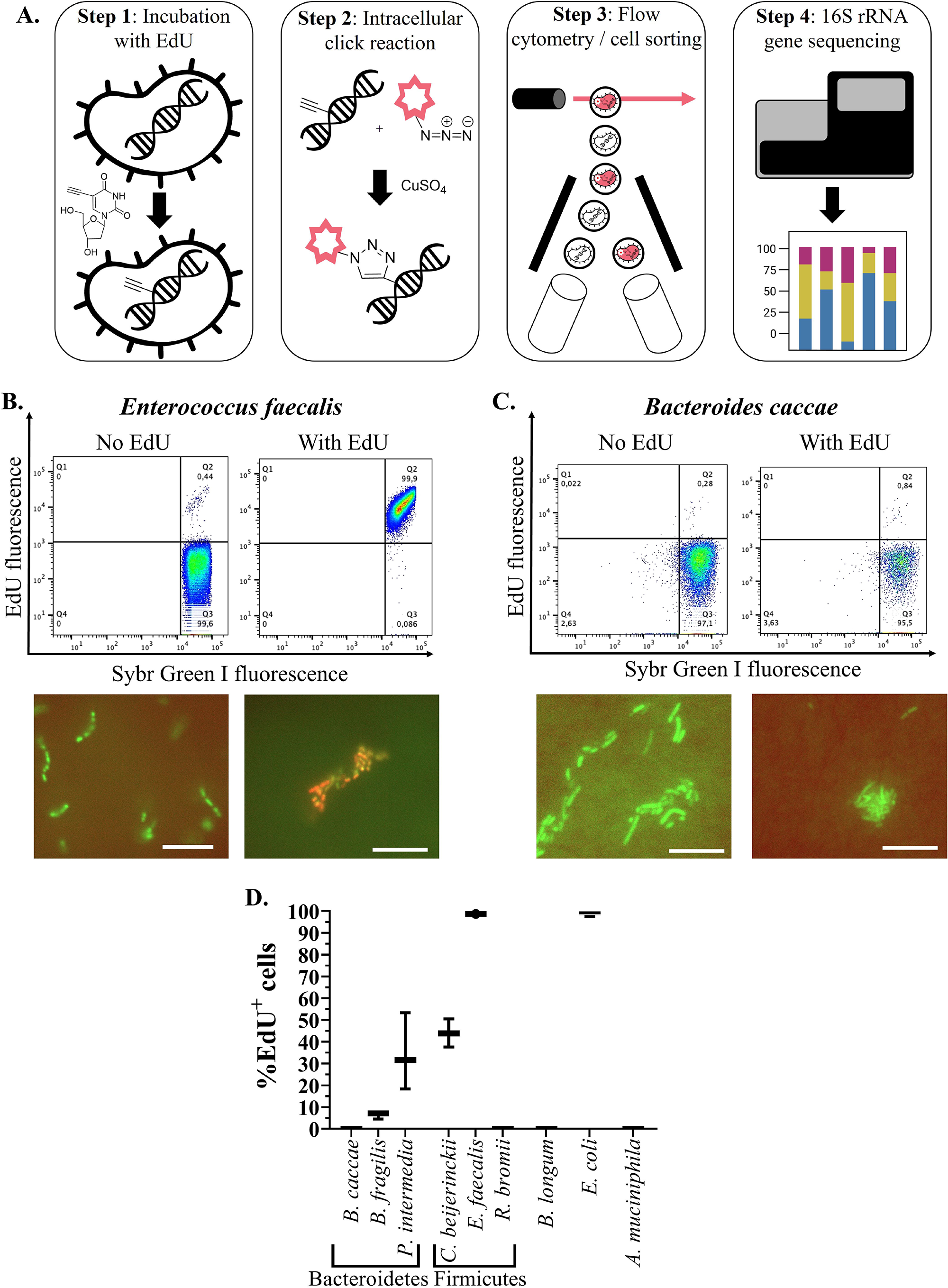
Optimization of EdU-click in gut bacterial isolates. Gut bacterial isolates underwent the EdU-click procedure as described in the main text. (A) Overview of the EdU-click and FACS-Seq procedure. In Step 1, bacterial cells are incubated *in vitro* with EdU. In Step 2, the alkyne group of the EdU molecule, located within the bacterial DNA, is exposed to a fluorescent azide and undergoes a copper-catalyzed ‘click’ reaction. In Step 3, bacteria which have incorporated EdU become fluorescent, are detected, and physically separated from unlabeled bacteria. In Step 4, bacterial DNA from both labeled and unlabeled bacteria undergo 16S amplicon sequencing and taxonomic analysis. (B) Top: Representative flow cytograms of *Enterococcus faecalis* grown either with or without EdU. Isolates grown without EdU were used to setup the gating strategy and determine background noise or unspecific labeling. Quadrant 3 (Q3): unlabeled, or EdU-negative (EdU-), population. Quadrant 2 (Q2): EdU+ population. Bottom: epifluorescence microscopy images of the same isolate in the flow cytograms above. Scale bar: 10 microns. (C) Top: Representative flow cytograms of *Bacteroides caccae* grown either with or without EdU. Bottom: epifluorescence microscopy images of the same isolate in the flow cytograms above. Scale bar: 10 microns. (D) Quantification of the proportion of EdU+ cells for each bacterial isolate. The proportion of EdU+ cells for each isolate was calculated as the average number of cells present in Q2, minus background noise, divided by the sum of cells in Q2 and Q3.

Bacterial isolates were incubated anaerobically at 37 °C with or without 20 µm EdU in reduced Bovine Heart Infusion broth (BHI) for three hours. Cells were then fixed and underwent the click and flow cytometry procedures as described in the Methods. Duplicates of each isolate incubated without EdU served as controls to set up the gating strategy (see **Figs 1b, c; S1**). The different bacterial isolates exhibited varying proportions of EdU^+^ cells, quantified relative to all bacterial cells obtained after SybrGreen staining (see Methods). Specifically, 34.38% ± 17.67 of *P. intermedia* cells, 43.97% ± 6.51 of *C. beijerinckii* cells, 98.7% ± 0.09 of *E. faecalis* cells, and 98.66% ± 0.31 of *E. coli* cells were labeled with EdU, whereas no cells of *B. fragilis*, *B. caccae, R. bromii, B. infantis,* or *A. muciniphila* were labeled during the same growth period (**Fig. 1b, d; Fig. S1a**). Heat-killed *E. coli* K12 cells which underwent the EdU-click procedure did not label (**Fig. S2**), demonstrating that only actively replicating bacteria are labelled using this technique.

It is unclear why some isolates did not label with EdU, despite exhibiting growth (**Fig. S1b**). It is not likely due to differences in cell wall type, since both Gram-positive (*E. faecalis, C. beijerinckii*) and Gram-negative (*E. coli, P. intermedia*) representatives were similarly labeled with EdU. Differences in growth rates may be implicated, as correlation analyses between the proportion of EdU^+^ cells and the maximal replicate rate (Max V) revealed a significant positive correlation (Pearson r^2^ = 0.407, p = 0.0005) for these isolates (**Fig. S1c**). This correlation was stronger when *B. infantis*, the only fast-growing isolate unlabeled with EdU, was removed from the analysis (r^2^ = 0.6019, p < 0.0001, **Fig. S1d**). Other possible reasons for the lack of labeling of these isolates are explored in more detail in the Discussion.

We further confirmed the fluorescence signals of these isolates by epifluorescence microscopy (**Fig 1b, c; Fig. S3**), validating the detected signal obtained by flow cytometry. Previous studies in *E. coli* have verified the specificity of EdU in labeling newly created DNA, and no other cell components.^31, 32^

### EdU-click validation in mouse fecal bacteria

We next sought to determine if EdU-click could be used to identify replicating bacteria in a complex microbial community, using fecal samples from healthy C57BL/6 mice.

We first tested incremental EdU concentrations from 10 to 100 µM on resuspended bacterial communities from mouse fecal samples, or mouse fecal bacteria, in three independent experiments (N= 4-5 mice per experiment, see Methods; **Fig 2a, Fig. S4**). The proportion of EdU^+^ cells varied according to the cohort of mice, limiting the direct comparison of proportions of EdU^+^ cells between experiments. In each independent experiment, the proportions of EdU^+^ cells increased in a concentration-dependent manner and were only significantly lower at 10 µM EdU relative to higher concentrations in one experiment (p < 0.05, Kruskal-Wallis with post-hoc Dunn’s multiple comparisons test, **Fig. S4a**). We report a non-significant increase in EdU^+^ cells with EdU concentrations between 10-35 µM (**Fig. 2a**), and between 20-80 µM (**Fig. S4b**). Since there were no large differences between these concentrations, we used the intermediate concentration of 20uM EdU for all downstream experiments.

Next, we tested two incubation media for EdU labeling of complex communities – reduced Bovine Heart Infusion broth (rBHI) and a fecal slurry. Aiming to best recapitulate the host environment, we performed EdU-click using each of these media either by themselves (100% rBHI or 100% fecal slurry) or as various ratios of the two (3:1, 1:1, and 1:3 ratio of rBHI to fecal slurry), (**Fig. 2b**). Incubation in 100% rBHI resulted in the largest proportion of EdU^+^ cells, decreasing with decreasing ratios of rBHI, while incubation in pure fecal slurry did not result in any EdU labeling, suggesting the slurry cannot support bacterial replication. The incubation conditions which contained the most fecal slurry and similar proportions of EdU^+^ cells to the rBHI were the 1:1 rBHI-fecal slurry medium, which is the one we selected going forward.

Finally, we determined the optimal incubation time with EdU. To minimize bacterial selection during incubation, we chose to explore incubation times that were 3 hours or less^29, 33^. We aimed to determine the shortest incubation time possible which still resulted in adequate labelling for downstream analyses (e.g., fluorescence-activated cell sorting). However, shorter incubation times resulted in significantly lower and less clear EdU^+^ signals, despite similar bacterial abundances (**Fig. 2c**). We therefore proceeded with 3-hour incubations going forward.

We then used the optimized procedure, being a 3-hour anaerobic incubation with 20 uM EdU in 1:1 rBHI-fecal slurry, to label the replicating fecal bacteria in a separate cohort of mice. As seen in the representative flow cytogram in **Figure 2d**, we obtained a clear population of bacterial cells distinct enough to proceed with cell sorting and sequencing of the replicating cells.

**Fig. 2.**
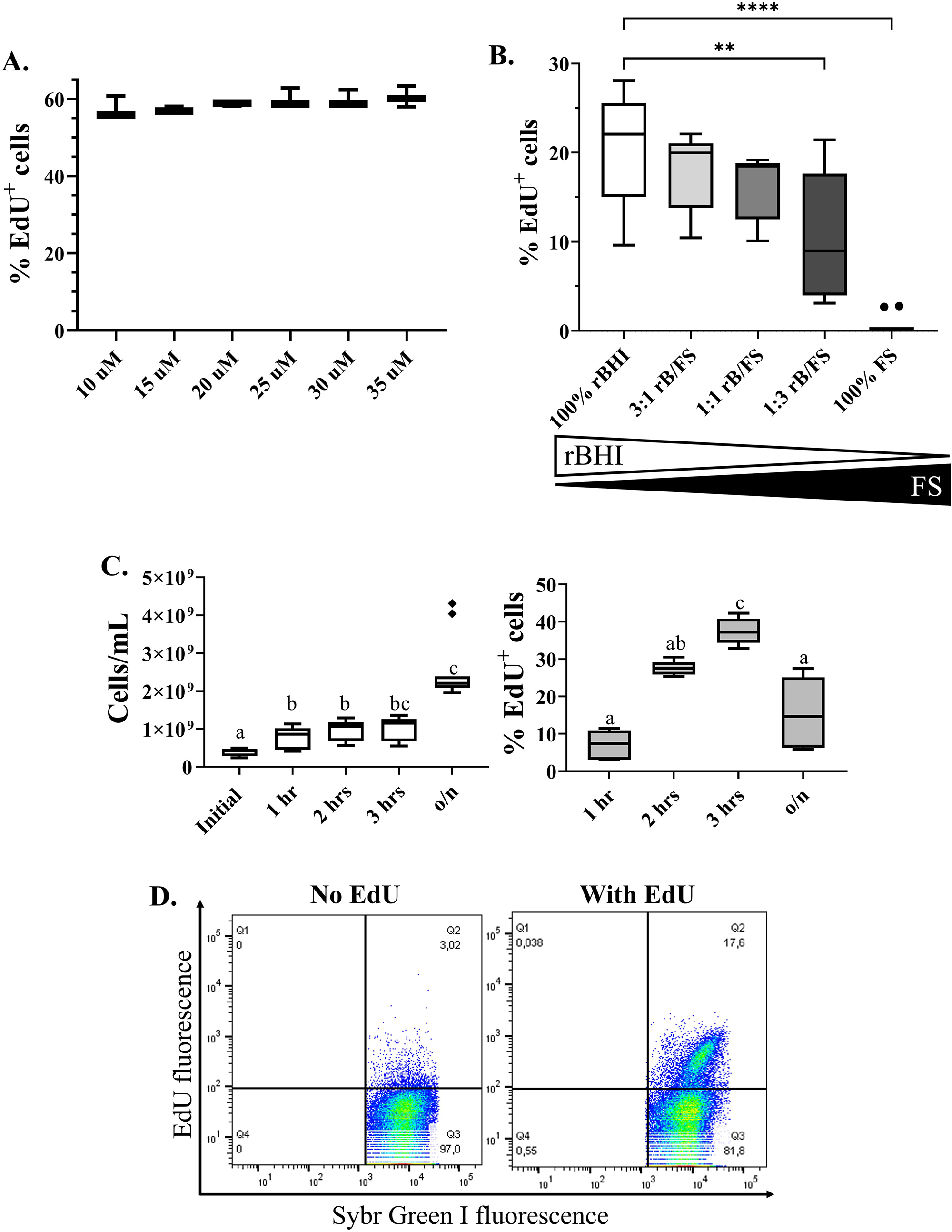
Optimization of EdU-click in fecal mouse bacteria. Mouse fecal bacteria underwent variations of the EdU-click procedure as described in the main text. (A) The proportion of fecal bacterial cells labeled as EdU+ using increasing concentrations of EdU. (B) The proportion of fecal bacterial cells labeled as EdU+ when using various combinations of media (rBHI) and fecal slurry. Comparisons are relative to 100% rBHI. Statistical significance determined by Kruskal-Wallis with Dunn’s test to correct for multiple comparisons. Comparisons are considered significant at p ≤ 0.05. rBHI = reduced Bovine Heart Infusion broth with hemin; FS = fecal slurry; rB/FS = rBHI and fecal slurry. (C) Left: The number of fecal bacterial cells, as calculated from the events during flow cytometry, after each incubation time. Statistical significance determined using Kruskal-Wallis with multiple comparisons corrected by Dunn’s test. Initial = no incubation; o/n= overnight (19 hours). Right: The proportion of fecal bacterial cells labeled as EdU+ after each incubation time. Statistical significance determined using Brown-Forsythe and Welch ANOVA with Dunnett’s multiple comparisons test Different letters denote a significant difference at p ≤ 0.05. (D) Representative flow cytogram of a mouse fecal bacterial community which underwent the optimized EdU-click procedure, incubated with or without EdU.

### EdU-click captures the actively replicating gut bacteria in healthy mice

Having optimized the EdU-click protocol for mouse fecal bacteria, we next used this technique to taxonomically identify the EdU^+^ fraction of cells in the feces of two cohorts of mice. Mouse fecal samples were obtained from a total of four cages of 4-5 mice per cage, encompassing both sexes, during two independent experiments (n=17). Fecal samples from all mice in each experiment were pooled before performing EdU-click to ensure enough original material for cell sorting. We then used fluorescence-activated cell sorting (FACS) to separate the following bacterial fractions: (1) the whole, unincubated initial community (initial community); (2) the EdU^+^ cells; (3) the EdU^-^ cells; and (4) the whole, incubated community (whole community).

Taxonomic identification of these sorted fractions, as well as the unsorted initial community, were conducted using 16S rRNA amplicon sequencing, as described in the Methods. The unsorted initial, sorted initial, and sorted whole community samples allowed us to determine possible cell sorting or incubation biases. Sequences present in the sorted communities and the sheath fluid used during cell sorting, but absent from the unsorted initial community, were identified as contamination and manually removed from downstream analyses (see Methods). The average sequencing depth of all samples (excluding the sheath fluid sample) was 24.04 Mbp (± 18.69 Mbp), the average number of reads per sample was 39,926 (± 31,039), and the rarefaction curve on the number of observed features found in each sample type is displayed in **Figure S5a**. After quality control and filtering of contamination, the average number of reads per sample was 20,614 (± 14,561), with samples keeping on average 54.64% (± 14.20) of their original reads.

The taxonomy at the genus level of the initial community and each sorted fraction (EdU^+^, EdU^-^, whole community) for the two experiments can be seen in **Figure 3a**. Notably, the taxonomy of the initial community consisted of more species of lower-abundant bacteria as compared to the sorted fractions, represented by the higher proportions of bacteria categorized as “Other”.

**Fig. 3.**
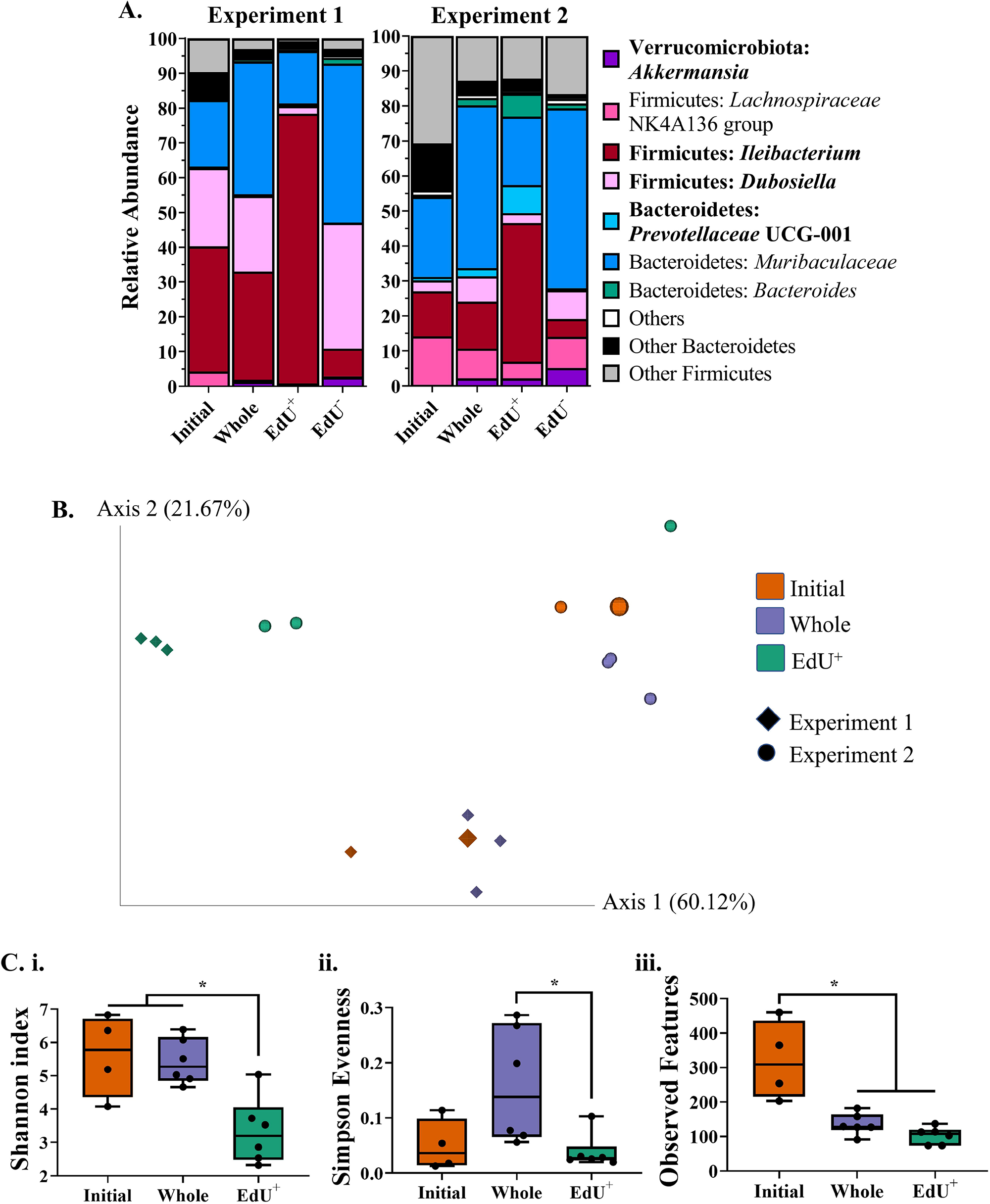
Taxonomy and diversity of fecal mouse bacteria after EdU-click. Mouse fecal bacteria from C57BL/6 mice (N= 4-5/ experiment) underwent the optimized EdU-click procedure and were subsequently sorted by fluorescence and analyzed by 16S amplicon sequencing. (A) The relative abundance of fecal bacteria for each experiment and each sorted fraction (initial and whole community, EdU+, and EdU-cells) at the genus level. The names of bacteria which are differentially abundant between the sorted fractions are bolded. (B) Principal coordinate analysis (PCoA) plot of weighted UniFrac distances for each sorted fraction in (A). The different shapes correspond to the two independent experiments. For the initial community, the larger point corresponds to the unsorted sample whereas the smaller point corresponds to the sorted sample. (C) Within-sample alpha diversity calculated using Shannon’s index (i), Simpson Evenness (ii), and Observed Features (iii) for each sorted fraction, combining the results from both independent experiments (N=17). Initial = initial community; Whole = whole community

Differential abundance analyses on the combined taxonomic data of both independent experiments identified which bacteria were more represented in the different sorted fractions, as described in the Methods. Some of the most important differentially abundant genera were, in order of importance, *Akkermansia, Prevotellaceae* UCG-001, *Ileibacterium,* and *Dubosiella*.

*Akkermansia* and *Dubosiella* followed similar patterns, as they were both depleted in the EdU^+^ sorted fraction despite increasing in relative abundance in the whole community. Indeed, *Akkermansia* had low relative abundances in the EdU^+^ fraction (experiment 1: 0.55% ± 0.21; experiment 2: 2.17% ± 2.82) and the Initial community (experiment 1: 0.03% ± 0.00; experiment 2: 0.28% ± 0.01) relative to the Whole community (experiment 1: 1.29% ± 0.38; experiment 2: 2.16% ± 0.30) and the EdU^-^ fraction (experiment 1: 2.61% ± 0.98; experiment 2: 5.16% ± 1.73). For its part, *Dubosiella* was decreased in the EdU^+^ fraction (experiment 1: 2.20% ± 0.04; experiment 2: 2.83% ± 3.95) as compared to the Whole community (experiment 1: 21.76% ± 6.27; experiment 2: 7.25% ± 2.14) and the EdU^-^ fraction (experiment 1: 36.24% ± 3.41; experiment 2: 8.23% ± 3.77). The increased relative abundance of the taxa which are not enriched in the actively replicating sorted fraction could result from the selective loss of other bacterial taxa, or could indicate that only a fraction of these cells are replicating to sustain this increase. Alternatively, it could suggest that despite actively replicating, they are not labeled by EdU-click, as seen in the human gut isolate of *A. muciniphila* in **Figure 1d**. The *Dubosiella* species contrasts with *Akkermansia*, however, in that it was already quite abundant in the initial community of experiment 1 (22.54% ± 1.65) but not in experiment 2 (3.13% ± 0.06). This species may have behaved differently in these two experiments due to the compositional differences of the initial input community (**Fig. 2a**).

The relative abundance of *Prevotella* UCG-001 was increased in only the EdU^+^ fraction (experiment 1: 0.44% ± 0.23; experiment 2: 7.96% ± 5.83) as compared to all of the other fractions, suggesting that despite being a relatively low-abundant bacteria – being present in the whole community with an average relative abundance of 0.44% ± 0.23 in experiment 1 and 2.33% ± 0.75 in experiment 2 – many of its members are actively replicating.

Similarly to *Prevotella* UCG-001, the relative abundance of *Ileibacterium* was increased in the EdU^+^ fraction (experiment 1: 77.59% ± 2.02; experiment 2: 39.66% ± 28.72) as compared to all of the other fractions. In contrast to *Prevotella* UCG-001, however, this taxon is already fairly abundant in the whole community (experiment 1: 31.09% ± 1.66; experiment 2: 13.42% ± 1.27).

The beta diversity of these bacterial communities was visualized using Principal Coordinate Analysis (PCoA) on the weighted Unifrac distance (**Fig 3b**). We excluded the EdU-sorted fraction from this and subsequent analyses since it resembled the whole community taxonomically (**Fig. 3a**). We report a significant difference in the weighted Unifrac distances between the two experiments (PERMANOVA, q = 0.008), as well as between the EdU^+^ fraction and the whole community (q = 0.018). These data support the fact that the EdU^+^ fraction is enriched in specific gut bacterial members.

When looking at the within-sample (alpha) diversity, the EdU^+^ fraction had significantly lower Shannon index values (Kruskal-Wallis, q = 0.029) in comparison to the other fractions for both experiments (**Fig. 3ci**). The EdU^+^ fraction also had a significantly lower Simpson Evenness (Kruskal-Wallis, q = 0.049) relative to the whole community (**Fig. 3cii**), suggesting the numerical dominance of certain taxa over others in the EdU^+^ fraction. Both of these findings are corroborated by the taxonomic composition of this fraction, which was enriched in certain genera and depleted in others, as described above (**Fig. 3a**).

The initial community had a significantly higher number of observed features (Kruskal-Wallis, q = 0.015) as compared to the other incubated fractions in both experiments (**Fig. 3ciii**), confirming that there are a greater number of species present in the initial community relative to the incubated fractions (**Fig. 3a**). The initial community also had a low Simpson evenness score (0.05 ± 0.047) (**Fig 3cii**), suggesting the abundance of some species over others. This is also in agreement with the taxonomy, as this fraction was typically composed of four main genera – *Muribaculaceae* (experiment 1: 19.09% ± 6.02; experiment 2: 22.94% ± 4.52), *Dubosiella* (experiment 1: 22.54% ± 1.65; experiment 2: 3.13% ± 0.06), *Ileibacterium* (experiment 1: 36.04% ± 6.13; experiment 2: 12.87% ± 1.28), and *Lachnospiraceae* NK4A136 group (experiment 1: 4.19% ± 1.44; experiment 2: 13.90% ± 4.36) (**Fig 3a**).

It is important to note that the Shannon index and Simpson Evenness metrics were significantly different between the two experiments (**Fig. S6a, b**), but the number of observed features was comparable (**Fig. S6c**). This is consistent with the fact that these experiments involved different cages of mice from different litters, factors known to influence gut bacterial community composition.^34^

Altogether, the diversity and taxonomic information obtained in this experiment strongly suggests that the EdU^+^ fraction is a specific subset of the whole gut bacterial community, enriched in *Prevotella* UCG-001 and *Ileibacterium*, and depleted in *Akkermansia* and *Dubosiella*.

### EdU-click reveals the replicating gut bacteria after an in vitro exposure to antibiotics

After identifying the replicating bacterial taxa in healthy mice, we next sought to determine how these populations would change upon an antibiotic perturbation, thereby highlighting the potential of this approach *in vitro*. We thus tested the effects of polymyxin B, an antibiotic targeting Gram-negative bacteria, and vancomycin, an antibiotic targeting Gram-positive bacteria, on bacteria from mouse fecal samples.

Fecal bacteria from healthy mice were incubated in the presence of either antibiotic or only the vehicle (DMSO), in addition to EdU where appropriate (see Methods). Sorting and sequencing were conducted on the same fractions as previously described. The average sequencing depth of all samples (excluding the sheath fluid sample) was 17.87± 12.21 Mbp, the average number of reads per sample was 29,685± 10,281, and the rarefaction curve on the number of observed features found in each sample type is displayed in **Figure S5b**. After quality control and filtering of contamination, the average number of reads per sample was 15,495 (± 9,164), with samples keeping on average 72.44% (± 14.40) of their original reads.

The growth curve in **Figure 4a** shows the impact of each antibiotic on the growth of mouse fecal bacterial communities, and **Table S1** quantifies several characteristics of these growth curves. Notably, polymyxin B resulted in a significantly longer lag time (One-Way ANOVA, p = 0.0077), slower replication rate (p = 0.0260), and less overall growth (p = 0.0013) as compared to the control (**Table S1b**). In contrast, vancomycin treatment only significantly affected total bacterial growth (p = 0.0455) and none of the other growth parameters measured (**Table S1b**).

The taxonomy of the initial community, as well as the taxonomy of each sorted fraction (EdU^+^, whole community) for each treatment (Control, Polymyxin B, Vancomycin) is displayed at the genus level in **Figure 4b**. When comparing the control to the Polymyxin B treatment, and accounting for sorted fraction as a random effect (see Methods), differential abundance analysis identified *Muribaculaceae* and *Akkermansia* as significantly decreased in the Polymyxin B treatment (*Muribaculaceae*: from 39.08% ± 15.58 to 5.51% ± 7.07; *Akkermansia*: from 3.17% ± 2.19 to 1.31% ± 1.72). This is consistent with the expected effect of this antibiotic on Gram negative bacteria. There were no differentially abundant species between the control and the Vancomycin treatment.

Differential abundance analysis on the sorted fractions for all treatments and accounting for treatment as a random effect (see Methods) revealed two differentially abundant genera: *Desulfovibrio* and *Ileibacterium*.

*Desulfovibrio* was differentially abundant due to its exclusion from the EdU^+^ group, as despite its low abundance in the other fractions (Whole: 1.04% ± 0.74; Initial 0.37% ± 1.15), it was virtually absent from the EdU^+^ fraction (0.03% ± 0.07). This suggests either that this bacterium is not actively replicating, or that it is not amenable to labeling with EdU.

As previously observed, *Ileibacterium* dominated the EdU^+^ fraction (67.08% ± 33.27) in comparison to the Whole (22.14% ± 11.07) and Initial (11.93% ± 1.14) fractions, regardless of the treatment. This contrasts with the known effects of vancomycin on Gram positive bacteria.

The PCoA on the weighted Unifrac distances for these conditions can be seen in **Figure 4c**. The EdU^+^ fractions of all the treatment groups cluster together and are distinct from the rest of the populations, and the EdU^+^ fraction is significantly different from the whole community (q = 0.045). There was no significant difference in the weighted Unifrac distance between any of the treatments (q > 0.05), again likely due to the significant clustering together of the EdU^+^ fractions from all treatment groups (**Fig. 4c**).

**Fig. 4.**
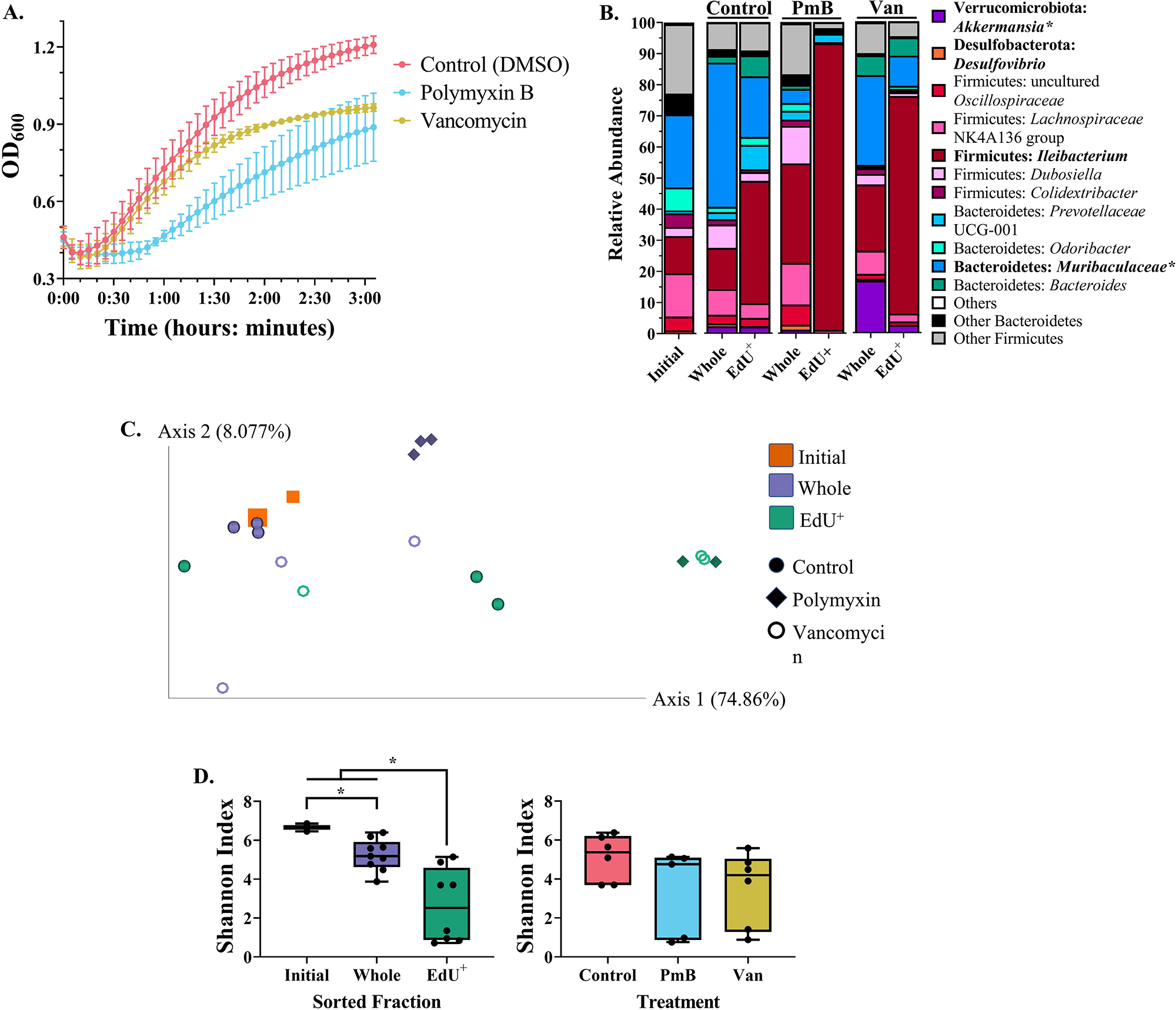
Identifying the replicating bacterial community during an in vitro antibiotic perturbation. Mouse fecal bacteria from two different pairs of cages (n = 4-5 mice per cage) were incubated in triplicate with polymyxin B, vancomycin, or no antibiotic (with DMSO, the drug solvent), as well as with or without EdU. The resulting communities underwent EdU-click, cell sorting, and 16S amplicon sequencing. (A) Growth curve of the fecal bacteria after addition of the antibiotic or solvent, where appropriate. (B) Taxonomic composition of the initial and whole community, and the EdU+ cells, for each treatment group at the genus level. Both the names of the phylum and genus are provided as Phylum: Genus. Gram + bacteria are colored in shades of red and pink whereas gram – bacteria are colored in shades of blue, green, orange, and purple. The names of bacteria which are differentially abundant are bolded. Bacteria which were differentially abundant between the different treatments have an asterisk (*) next to their names, whereas differentially abundant bacteria between the sorted fractions do not. (C) Between-group beta diversity (weighted UniFrac), colored by sorted fraction and with different shapes for each treatment group. For the initial community (squares), the larger point corresponds to the unsorted sample whereas the smaller point corresponds to the sorted sample. (D) Left: Within-group alpha diversity (Shannon’s index) of each sorted fraction, regardless of treatment. Right: Within-group alpha diversity (Shannon’s Index) of each treatment group, regardless of sorted fraction. Differences in alpha diversity were determined by pairwise PERMANOVA. Significance is denoted as q ≤ 0.05. PmB = polymyxin B; Van = vancomycin.

The Shannon index values for each treatment condition can be seen in **Figure 4d**. Once again, the EdU^+^ fractions had a significantly lower Shannon index as compared to the whole community (q = 0.016) and the initial community (q = 0.037) sorted fractions, regardless of treatment type (exposure or not to either antibiotic). Furthermore, the initial community had a significantly higher Shannon index as compared to the whole community (q = 0.037). When considering the different treatment types regardless of the sorted fraction, there were no significant differences in the Shannon index values between the treatments, possibly due to the low Shannon index values in all EdU^+^ fractions (**Fig. 4d**).

## Discussion

We describe here the optimization of EdU “click” chemistry (EdU-click) paired with FACS and 16S amplicon sequencing to identify replicating gut bacteria *in vitro*. We first show that EdU-click labels gut bacterial isolates with no bias based on cell wall type, yet reveals notably fewer Bacteroidetes labeled relative to Firmicutes (**Fig. 1d**). The results we have gathered thus far suggest genus-level variation in EdU labeling, though more thorough investigations with other gut bacterial isolates are needed to confirm this.

We note that not all gut bacterial isolates were amenable to EdU-click. As mentioned in the Results, this did not seem to be due to differences in cell wall type. There was, however, a significant positive correlation between the replication rate of an isolate and the proportion of its cells which were labeled (**Fig. S1c**), suggesting that bacteria must grow rapidly during the incubation with EdU in order to be labeled. However, replication rate alone cannot fully explain the discrepancies in labeling seen here, since *B. infantis* grew rapidly (**Fig. S1b**) but did not label (**Fig. S1a**).

Another possible explanation could be phylogenetic differences in the ability to uptake or use exogenous pyrimidine nucleosides (such as thymidine, of which EdU is an analogue), which has been shown to affect bacterial DNA labeling in other complex communities.^35, 36^ Even when bacteria encode the genes to use thymidine, some may preferentially use the *de novo* pathway rather than the salvage pathway to make nucleotides,^37^ the latter being required for EdU labeling. Further work on other gut bacterial isolates, especially members of the phyla with many unlabeled representatives, is thus required to identify the underlying mechanisms.

Regardless of the reason for the lack of labeling of certain bacterial isolates, these results confirm that EdU-click can only inform about the replication dynamics of bacteria which are both amenable to labeling and which are actively replicating. This is a known problem when using thymidine analogues to track replicating bacterial populations, as mentioned previously.^35, 36, 38^. Since the ability of all environmental microorganisms to uptake thymidine analogues like EdU remains undetermined, the results from EdU-click should be interpreted cautiously when used in microbial assemblages.

Despite these inherent limitations, we optimized the EdU-click procedure for mouse fecal bacterial communities, as even partial knowledge of the replicating gut bacteria could provide a more nuanced understanding of gut microbial dynamics, community metabolism and, ultimately, host metabolism.^15, 21^ Based on our experience, the EdU-click optimization process we have done here should be done for any other microbial ecosystem of interest, following our framework.

When characterizing the EdU^+^ cells derived from the feces of healthy mice, we report that the EdU^+^ fraction had significantly lower alpha diversity metrics, and a significantly different weighted Unifrac distance, as compared to the whole community, regardless of the cage of origin. These results suggest that the EdU^+^ fraction is a specific subset of the whole community. This is further supported by taxonomic analysis, which revealed that the EdU^+^ fraction was enriched in *Ileibacterium* and *Prevotella* UCG-001, and depleted in *Akkermansia* and *Dubosiella*, as compared to the whole community.

There is not much literature on *Ileibacterium*, with only one characterized species, *I. valens*, in 2017.^39^ So far, *I. valens* has been seen to increase in abundance after dietary changes,^39–42^ and it has been associated with a few immunological and host physiological parameters.^42^ Our results here suggest that it may be a very actively replicating member of the mouse gut microbiota, though its specific roles in the gut are yet to be determined.

The EdU procedure developed here requires an incubation step which can still lead to an altered community structure, despite our efforts to minimize this. Indeed, the diversity and taxonomic composition of the initial community differed from that of the incubated whole community in all experiments in this study. Namely, incubation resulted in decreased richness due to the loss of less abundant taxa, either not supported by the incubation medium or lost during the click procedure and sorting steps (**Fig. 3b,e; Fig. 4b,d**).

EdU-click may thus capture the replication potential of the more abundant gut bacterial members which are amenable to labeling, rather than accurate *in situ* replication of all members present. Nevertheless, replication potential has been shown to strongly correlate with *in situ* replication,^43^ and *in vitro* incubations of microbial communities for several hours outside their original ecosystems have been shown to not alter the original *in situ* replication patterns.^33^ Overall, this suggests that EdU-click can still provide useful information on *in situ* dynamics for the most abundant bacterial members or those with a higher growth rate, despite necessitating an *in vitro* incubation step.

Finally, we demonstrate one potential application of our optimized EdU-click technique in an *in vitro* antibiotic perturbation experiment on mouse fecal bacteria incubated with either polymyxin B or vancomycin. Polymyxin B exposure resulted in a decrease in the relative abundance of *Muribaculaceae* and *Akkermansia* (both Gram-negative bacteria), whereas vancomycin did not have a large effect on the bacterial community or its Gram-positive members. Importantly, *Ileibacterium* constituted the vast majority of the EdU^+^ fraction in both the polymyxin B and the vancomycin treatment.

One explanation for the replication of *Ileibacterium* during exposure to vancomycin is that this taxon was replicating and incorporating EdU before the antibiotic affected its growth. This is supported by the growth curve of the vancomycin-treated bacterial community, wherein the bacteria were replicating at similar rates as the control community until later in the growth curve (**Fig. 4a, Table S1**). We cannot rule out other possibilities, however, such as insufficient dosage of the antibiotic (despite using a final concentration of 0.1 mg.mL^-1^, higher than the estimated vancomycin minimal inhibitory concentration [MIC] of 0.001-0.01 mg.mL^-1^ for these samples; data not shown) or innate resistance of *Ileibacterium* to vancomycin.

To investigate the latter, we identified genes in the only sequenced *I. valens* genome available which were homologous to known antimicrobial resistance genes, using the Comprehensive Antibiotic Resistance Database (CARD) Resistance Gene Identifier (RGI) on the translated protein sequences of this genome.^39, 44^ The results were inconclusive, as there were only loose hits to a variety of antibiotic resistance genes. These included hits to 21 vancomycin and glycopeptide resistance genes (**Table S2**) as well as 69 hits to genes related to various antibiotic efflux pumps (data not shown). Despite being loose hits, either strategy – glycopeptide resistance or antibiotic efflux – could explain the ability of *Ileibacterium* to replicate during exposure to vancomycin. Furthermore, other members of the *Erysipelotrichaceae* family, to which *Ileibacterium* belongs, have been reported as being intrinsically resistant to vancomycin.^45–47^ Considering that mechanisms for resistance against glycopeptide antibiotics like vancomycin can differ greatly between taxa and remain to be fully characterized in all bacteria,^48^ it is thus still possible that *Ileibacterium* is truly resistant to vancomycin.

Overall, we show that EdU-click enables the rapid detection of replicating bacteria, both in clonal populations and in heterogenous microbial assemblages. The procedure is not laborious and it gives valuable qualitative information on bacterial replication. Furthermore, complementing EdU-click with cell sorting and DNA sequencing allows for the taxonomic characterization of actively replicating bacteria. Coupling this information with knowledge of the taxonomic composition of the overall bacterial community can help reveal differential bacterial replication strategies in homeostatic and disrupted conditions, as observed in this work.

Our modified EdU-click procedure can best be applied for studying bacterial replication *in situ* and *in vitro*, where the ability of the resident bacteria to be labeled with EdU has been determined beforehand. For instance, bacterial communities known to label with EdU could be incubated with EdU in their natural environment (e.g., seawater, soil slurries, activated sludge). This technique would also be useful for tracking the replicating members of simplified bacterial communities in artificial gut systems (e.g., intestinal organoids, bioreactors, chemostats), or when only the replication of specific taxa is of interest. Furthermore, tracking replicating bacterial communities over time using EdU-click will help reveal the dynamics of specific bacterial replication and their impacts on the metabolism and composition of bacterial communities. This technique will be especially informative for tracking changes in bacterial replication and composition during the establishment of bacterial communities (e.g., in early life), during homeostasis, and as changes occur in the environment of these bacteria (e.g., antibiotic administration, intestinal pathogen infection).

Going forward, it would be worth further investigating the cause(s) of the differential labeling of bacterial isolates with EdU and quantifying the amount of EdU entering the cells. Importantly, uncovering the possible roles of *I. valens* in the colons of healthy mice and its apparent resistance to vancomycin will also be key to better understand the resilience of certain bacterial taxa to perturbations.

## Methods

### Animals

Feces from male and female wild-type C57BL/6 mice (Jackson Laboratories) kept at the Goodman Cancer Center at McGill University were collected, in accordance with the McGill Ethics Research Board (animal ethics protocol #5287). Over the course of the EdU optimization process and *in vitro* experiments, different cages of mice with varying ages were used. During the experiments on the healthy mouse gut microbiota, mice were matched for age. In experiment 1, the mice were 6 weeks and 6 days old and consisted of 4 male mice and 4 female mice of the same origin. In experiment 2, the mice were 7 weeks and 3 days old, and consisted of 4 male mice and 5 female mice of the same origin, with the except of one female mouse which was from a different litter. It should be noted that the same feces from the mice in experiment 2 were used for the antibiotic perturbation experiment (see *EdU-click performed on antibiotic-treated mouse fecal bacteria*).

In all cases, experiments were conducted on a mixture of feces from both male and female mice. Fecal pellets were collected from the mice by putting each mouse into its own cage, empty of corncob bedding and nesting material, with no access to food or water. Feces from each mouse were collected soon after defecation. Feces from mice of the same cage were put into the same 1.5 mL microcentrifuge tube. This procedure lasted no longer than one hour, after which mice were placed back into their respective cages. This procedure was never repeated more than twice in one week. Immediately after this collection procedure, the fecal pellets were either processed in an anaerobic chamber as detailed in *Isolation of bacteria from mouse* feces or frozen at -80°C.

### Bacterial strains

The bacterial strains used in this study were *Bacteroides fragilis* (32-6-I 11 MRS AN), *Escherichia coli K12 (*ATCC 25404*), Bifidobacterium longum* subsp. *infantis (*ATCC 15697*), Bacteroides caccae (*ATCC 43185*), Prevotella intermedia (*ATCC 25611*), Clostridium beijerinckii (*ATCC 51743*), Ruminococcus bromii (*ATCC 27255*), Akkermansia muciniphila (*ATCC BAA-835), and *Enterococcus faecalis* (SF24397). Each bacterium was originally isolated from the human gut, with the exception of *P. intermedia*, which comes from the human oral cavity. All bacteria were cultured anaerobically in an anerobic chamber (Coy Laboratory Products; 5% H2, 20% CO2, 75% N2) at 37°C overnight directly from frozen (-80°C) glycerol stocks, in autoclaved reduced Bovine Heart Infusion (rBHI) broth (Oxoid).

### Absorbance spectroscopy and EdU labeling of bacterial isolates

Overnight cultures of bacterial isolates were diluted in rBHI in a Falcon multiwell 24-well plate (Corning) to an optical density at 600 nm (OD600) of 0.05, with six replicate wells per isolate; three replicates were incubated with 20 μM EdU (from the Click-iT EdU Alexa Fluor^TM^ 647 flow cytometry assay kit, ThermoFisher Scientific) and three replicates were not incubated with EdU. For those replicates incubated without EdU, an equivalent volume of reduced 1X 0.2 μm-filtered phosphate-buffered saline (rPBS – 1 mg mL−1 L-cysteine) (Bioshop) was added to each well. For *E. faecalis* and *E. coli* K12, only two wells were used for the microscopy and replication measurements – one incubated with EdU, and one without. Additionally, *R. bromii* had two wells incubated with EdU and two without, due to the limited growth of the overnight culture.

All bacteria were incubated anaerobically in the 24-well plate at 37°C for 3 hours in an Epoch 2 Microplate Spectrophotometer (BioTek), with optical density measured at 600 nm (OD600). Every 15 minutes, the spectrophotometer shook the plate in a double-orbital fashion for 10 seconds and read the absorbance of each well. After the incubation, the bacteria were immediately fixed in an equal volume of 80% ethanol, for a final concentration of 40% ethanol. Fixed bacteria were either stored at 4°C overnight or processed after 10 minutes of incubation in the ethanol. The same or the next day, bacterial isolates underwent the click reaction as described in *Click reaction for bacterial isolates and fecal-derived bacteria*, and were either observed using flow cytometry as described in *Nucleic acid staining, flow cytometry, and fluorescence-activated cell sorting* or were prepared for microscopy as described in *Fluorescence microscopy of bacterial isolates*.

### Click reaction for bacterial isolates and fecal-derived bacteria

The following procedure is the optimized version of the EdU-click method. For both bacterial isolates and fecal-derived bacteria, fixed bacteria were pelleted by centrifugation (8,000xg, 5 min) and washed in 1X 0.2 μm-filtered phosphate-buffered saline (PBS) (Bioshop) before undergoing the click reaction as described by the kit protocol, with the exception that a final concentration of 5 μM of the Alexa Fluor^TM^ 647 azide fluorophore (purchased separately as Alexa Fluor^TM^ 647 azide, Triethylammonium Salt from ThermoFisher Scientific) was used. Afterwards, the bacteria were again pelleted by centrifugation (8,000xg, 5 min) and washed in an equal volume of 80% ethanol (final concentration 40%). The bacteria were resuspended in the diluted ethanol using filter pipettes, then they were centrifuged once more (8,000xg, 5 min), the ethanol was removed, and the pellet was resuspended in PBS for subsequent visualization using flow cytometry or fluorescence microscopy.

### Nucleic acid staining, flow cytometry, and fluorescence-activated cell sorting

After the EdU-click procedure, bacterial cells were diluted as needed (between 1:100 and 1:200) in PBS in 5 mL polystyrene round-bottom tubes (Falcon). All samples were stained with 1X Sybr Green I nucleic acid gel stain (Sybr Green I; Invitrogen by Thermo Fisher Scientific) for 15 minutes before acquisition on a flow cytometer. Flow cytometric measurements were obtained using the FACS Canto II (BD Biosciences, San Jose, CA, USA) equipped with a solid-state blue laser (488 nm, 20-mW) and a He/Ne ion red laser (633-nm, 17-mW) with BD FACSDiva software (BD Biosciences). The blue laser was used to excite Sybr Green I, and the green-fluorescence emission from Sybr Green was collected in the FITC channel (BP 530/30 filter). The red laser was used to excite the Alexa Fluor^TM^ 647 fluorophore, and the red-fluorescence emission from this fluorophore was collected in the APC channel (BP 660/20). Forward angle light scatter (FSC) was collected in the FSC channel, and side angle light scatter (SSC) was collected in the SSC channel (BP 488/10 filter).

When the bacterial cells underwent fluorescence-activated cell sorting (FACS), the FACSAria II (BD Biosciences, San Jose, CA, USA) was used, which is equipped with a solid-state blue laser (488 nm, 20-mW) and a He/Ne ion red laser (633-nm, 17-mW) with BD FACSDiva software (BD Biosciences). The same excitation and emission parameters as used during flow cytometry were also used for cell sorting.

Data obtained from flow cytometry and FACS were analyzed using FlowJo^TM^ software. Cell numbers were calculated from flow cytometry data using standards of known size and number. EdU+ cells were classified as those events which appeared in the upper right section of the flow cytogram (Q2). EdU-cells were classified as those events which appeared in the lower right section of the flow cytogram (Q3) when the sample was incubated with EdU. Any events which appeared in Q2 when the sample was incubated without EdU were classified as noise and subtracted from the number of EdU+ cells. The proportions of EdU+ cells were quantified as the number of cells in Q2 minus the average amount of noise and divided by the sum of cells present in Q2 and Q3.

### Fluorescence microscopy of bacterial isolates

After undergoing the click reaction as described in *Click reaction for bacterial isolates and fecal-derived bacteria*, bacterial isolates were prepared for microscopy in the following way. Bacterial isolates suspended in PBS were further diluted in PBS as necessary, and then portioned into 1 mL aliquots. These aliquots were stained with 1X Sybr Green I nucleic acid gel stain (Sybr Green I; Invitrogen by Thermo Fisher Scientific) for 15 minutes, and then these aliquots were vacuum filtered through 0.22 um filters on top of 0.3 um filters. Once the filters were dry, they were placed onto a microscope slide that had a drop of Citifluor AF1 Mountant Solution (Electron Microscopy Sciences) on it. A coverslip treated with this same antifade was then placed on top of the filter. The coverslip was sealed onto the microscope slide using commercially available nail polish. Once the slides were sealed, they were stored at -20°C until they were visualized.

Visualization of slides typically occurred the day after the slides were prepared, but never later than 1 week after slide preparation. All slides were visualized using the Axiovert 3 widefield inverted microscope (Zeiss) equipped with a Zeiss Axiocam 506 monochrome CCD camera (2752 x 2208 pixels, 4.54 um pixels) and Sola SM solid state light engine, using the 100X objective (PLAN-APOCHROMAT, NA=1.4, OIL, Iris) under oil immersion and the FS 24 (FITC) and FS 32 (Cy5) fluorescence cubes. Images were further processed in Fiji (version 1.53q).

### Isolation of bacteria from mouse feces

Fresh fecal pellets were collected in 1.5 mL microcentrifuge tubes from wildtype C57BL/6 mice on the day of EdU incubation. Fecal pellets were diluted as 1 μg of feces in 10 μl liquid (1:10) in 1.5 mL microcentrifuge tubes using reduced 1X 0.2 μm-filtered phosphate-buffered saline (rPBS – 1 mg mL−1 L-cysteine) (Bioshop) and vortexed thoroughly until no solid pellet remained (∼2-5 minutes). The tubes were then centrifuged for 1 minute at 700xg, and the supernatant was removed and added to a new 1.5 mL tube. These new tubes were centrifuged for 3 minutes at 6,000xg, and the resulting supernatant from all tubes was added into one 15 mL centrifuge tube. This supernatant, called the “fecal slurry”, was saved as part of the medium in which the bacteria would be incubated. The pellet was then resuspended in rPBS and centrifuged again, for 2 minutes at 6,000xg. The supernatant was discarded and the pellet was resuspended once more in rPBS. The resulting mixture was used as fecal-derived bacterial cells.

### EdU labeling of fecal-derived bacteria

The following procedure is the optimized version of the EdU-click method; note that the incubation medium, EdU concentration, and incubation time were different during optimization (see Results). Fecal-derived bacteria were diluted 1:10 in a medium made of 50% rBHI and 50% fecal slurry (1:1 rB/FS). These bacteria were incubated anaerobically at 37°C for 3 hours in 1.5 mL microcentrifuge tubes in triplicate or duplicate with or without a final concentration of 20 μM EdU, respectively. Immediately after the incubation, bacterial cells were fixed in an equal volume of 80% ethanol (final concentration 40%) and stored at 4°C overnight. The next day, fecal-derived bacteria underwent the click reaction and were observed using flow cytometry as described in *Click reaction for bacterial isolates and fecal-derived bacteria* and *Nucleic acid staining, flow cytometry and fluorescence-activated cell sorting*, respectively.

### EdU-click performed on feces of healthy mice

Fecal-derived bacteria were isolated from two cages of mice (one male cage, one female cage), as described under *Animals*, in two independent experiments. In each experiment, feces from each cage were pooled into their own 1.5 mL tube. EdU-click was performed as described in *EdU labeling of fecal-derived bacteria*, except that in each experiment, the bacteria were first incubated in 3 wells of a 96-well plate for 1 hour in an Epoch 2 Microplate Spectrophotometer (BioTek), with optical density measured at 600 nm (OD600). Every 15 minutes, the spectrophotometer shook the plate in a double-orbital fashion for 10 seconds and read the absorbance of each well. After this 1-hour incubation, 2 µl of DMSO was added to the culture in addition to 20 µM (final concentration) of EdU or an equal volume of rPBS, where appropriate, and the plate was incubated for another 3 hours in the spectrophotometer, after which the bacteria were fixed and stored as described in *EdU labeling of fecal-derived bacteria*.

After the click reaction and nucleic acid staining were performed as described in *Click reaction for bacterial isolates and fecal-derived bacteria*, bacterial cells underwent cell sorting as described in *Nucleic acid staining, flow cytometry and fluorescence-activated cell sorting*. The following populations were sorted: (1) Whole, unincubated community (initial community); (2) EdU+ cells; (3) EdU-cells; and (4) Whole, incubated community. Populations (1) and (4) were sorted based on their Sybr Green I nucleic acid signal, whereas populations (2) and (3) were sorted from each other based on their Alexa Fluor^TM^ 647 signal (for EdU incorporation).

### Antibiotics

The antibiotics used in this study were vancomycin (vancomycin hydrochloride from *Streptomyces orientalis*, Sigma Life Sciences lot # 085M4163V) and polymyxin B (polymyxin B sulfate, BioShop lot # 4F34412). Both were dissolved in 100% dimethyl sulfoxide (DMSO) to a concentration of 10 mg/mL and stored at 4°C protected from light. Dilutions of this stock were made and added to fecal-derived bacterial cultures at the appropriate time (see *EdU-click performed on antibiotic-treated mouse fecal bacteria*). The final concentration of both antibiotics was 1 µg/mL. This concentration was chosen based on our assessment of the minimum inhibitory concentration (MIC) for these antibiotics in the specific media we used (data not shown).

### EdU-click performed on antibiotic-treated mouse fecal bacteria

Bacteria were extracted from fresh mouse fecal pellets obtained from healthy, wildtype C57BL/6 mice (see *Animals*) as described under *Isolation of bacteria from mouse feces*. These bacteria were incubated anaerobically in an Epoch 2 Microplate Spectrophotometer (BioTek) at a 1:10 ratio in a final culture volume of 200 µl per well in 12 wells of a 96-well plate for 1 hour in 1:1 rBHI:fecal slurry. Every 15 minutes, the spectrophotometer shook the plate in a double-orbital fashion for 10 seconds and read the optical density of each well at 600 nm (OD600).

After this 1-hour incubation, the appropriate concentration of vancomycin, polymyxin B, or DMSO (the vehicle), was added to the culture in addition to 20 µM (final concentration) of EdU or an equivalent volume of rPBS, where appropriate. Each treatment – being vancomycin with EdU, polymyxin B with EdU, DMSO with EdU, and DMSO without EdU – had 3 wells dedicated to it (i.e., each treatment was performed in triplicate). After administration of the antibiotics, the plate was incubated for an additional 3 hours in the same conditions as the 1-hour incubation, after which the bacteria were fixed in an equal volume of 80% ethanol (final concentration 40% ethanol) and stored overnight at 4°C. The following day, the click procedure was performed on these samples as described under *Click reaction for bacterial isolates and fecal-derived bacteria*. The EdU+ and EdU-bacteria, as well as the initial and whole community, were then sorted from these samples using FACS, as described under *Nucleic acid staining, flow cytometry and fluorescence-activated cell sorting*. DNA extraction and 16S rRNA gene sequencing were performed on these populations as described under *DNA extraction from fecal-derived bacterial cells* and *16S rRNA amplicon sequencing and analysis*.

### DNA extraction from mouse fecal-derived bacterial cells

Bacterial DNA was extracted from sorted and unsorted mouse fecal-derived bacterial cells using the AllPrep PowerFecal DNA/RNA kit (Qiagen) as per manufacturer instructions, with the following alterations: 200 µl of sorted bacterial cells were used instead of 200 mg stool, all cold incubations were conducted in an ice bucket instead of at 4°C, the amount of supernatant and buffer C4 used after IRS incubation was doubled, the extracted DNA was diluted in RNAse free water rather than EB buffer, and the RNA collection steps were not performed. For the cell lysis step, bead-beating was performed using a Vortex Genie 2 with a microtube holder (SI-H524) (Scientific Industries) for 10 minutes when extracting <10 samples at a time, or for 20 minutes when extracting >10 samples at a time. All extracted DNA was immediately stored at -20°C.

### 16S rRNA gene amplicon sequencing and analysis

For both the experiment on the feces of healthy mice and the antibiotic *in vitro* perturbation experiment, extracted DNA underwent library preparation and 16S rRNA gene sequencing of the V4-V5 hypervariable region using the Illumina MiSeq PE250 system at the Université de Québec à Montréal (UQAM) Center of Excellence in Research on Orphan Diseases – Foundation Courtois (CERMO-FC) genomics platform.

The resulting bacterial sequences were analyzed using QIIME2 version 2022.2.^49^ As described in the Results, any sequences present in the sheath fluid but absent in the initial unsorted community were identified and manually removed from the analysis. Briefly, we used DADA2 in QIIME2 for quality control and filtering. Sequences were rarefied to a depth of 4458 reads for the healthy mice and 4702 reads for the antibiotic experiment before alpha and beta diversity analyses. Specifically, we used Shannon’s and Simpson’s indices for alpha diversity, and weighted Unifrac for beta diversity. Principal component analysis (PCoA) plots were constructed using weighted Unifrac distances. Significance between alpha diversity measures based on metadata (e.g., treatments, cages, sex, etc.) was measured using a Kruskal-Wallace test with pairwise comparisons. Significance between beta diversity measures was conducted using pairwise PERMANOVA comparisons with 999 permutations.

Taxonomic classification and relative abundance measurements were conducted on the quality-controlled, filtered sequences using QIIME2’s machine-learning program pre-trained on the 16S V4-V5 regions, using the Silva database.

Differential abundance analyses for all experiments were carried out in R (version 4.1.3) with ANCOM-II with Benjamini-Hochberg correction for multiple comparisons.^50^

### Statistical analyses

All statistical tests outside of those conducted in QIIME2 were conducted using Prism version 9.3.1 (GraphPad). Results were determined to be significantly different at P-value ≤ 0.05. Significance testing was conducted using the Kruskal-Wallis test with Dunn’s multiple comparisons test, Brown-Forsythe and Welch ANOVA with Dunnett’s multiple comparisons test, or one-way ANOVA with Dunnett’s test, as described in the Figure legends.

Correlation analyses between bacterial isolate maximal replication rates and their respective proportions of EdU+ cells were conducted by calculating Pearson’s correlation coefficients in Prism. Area under the curve (AUC) calculations of all bacterial growth curves was also calculated in Prism.

## Data Availability Statement

Bacterial 16S rRNA gene sequencing data can be accessed on the NCBI SRA database, accession number PRJNA855973. Code related to the analysis has been deposited in GitHub (https://github.com/evetb). Raw microscopy and flow cytometry data have been deposited in FigShare (https://doi.org/10.6084/m9.figshare.c.6103014.v1).

## Authors’ contributions

ETB designed, performed, analyzed, and interpreted experiments. CH helped with the bacterial isolate experiments. CFM helped develop the project, obtained funding, helped design experiments, and interpret the data. ETB and CFM edited the manuscript and approved the final draft.

## Acknowledgements

The authors would like to thank all the members of the Maurice lab, as well as Professor Bastien Castagner and Professor Gregory Marczynski, for their help editing parts of this manuscript. The flow cytometry work was performed in the McGill University Flow Cytometry Core Facility for flow cytometry and single cell analysis of the Life Science Complex and supported by funding from the Canadian Foundation for Innovation. The microscopy was performed in the McGill University Advance Bioimaging Facility (ABIF), RRID:SCR_017697.

## Figure Legends

**Figure S1.**
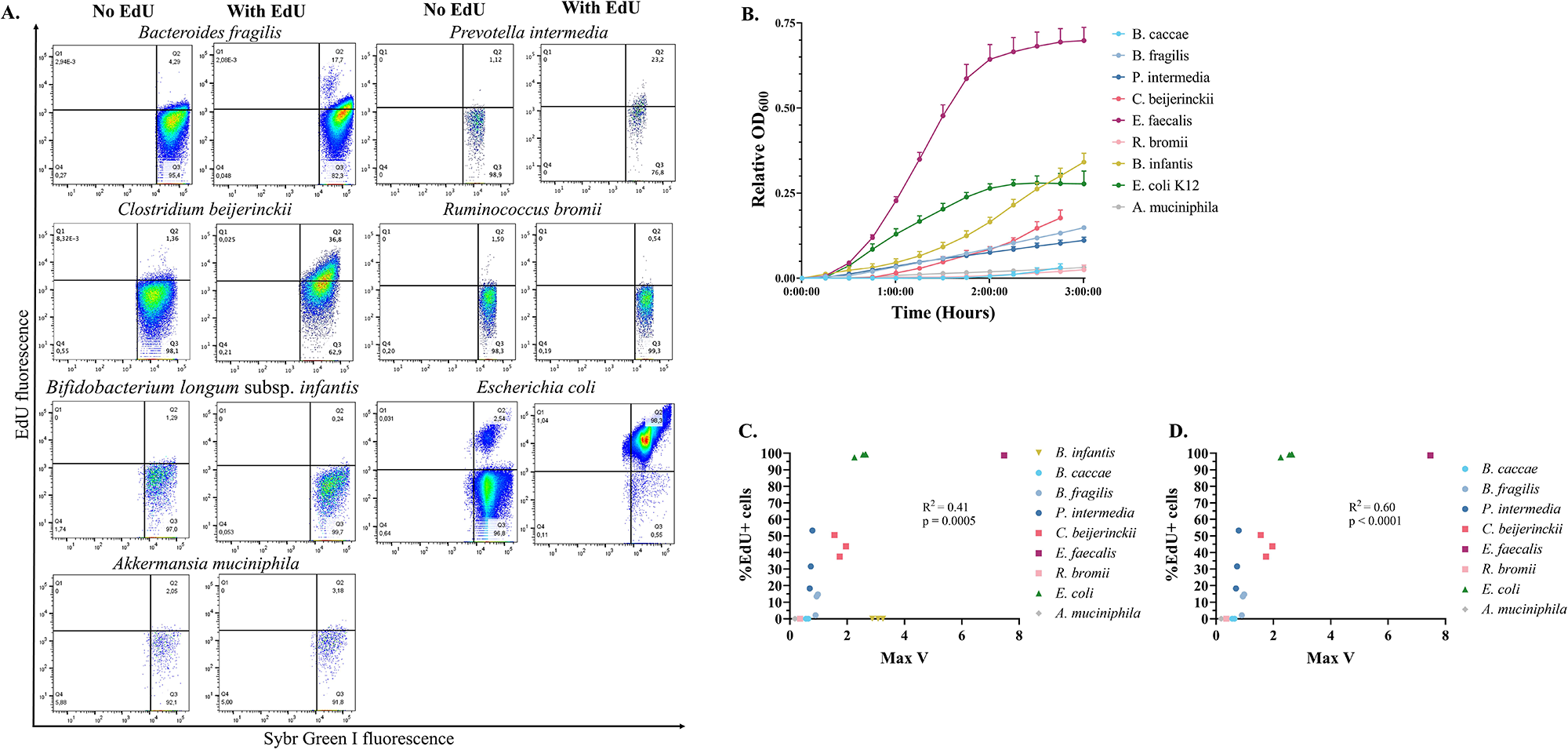
Variable labelling and growth of gut bacterial isolates with EdU. Gut bacterial isolates underwent a click reaction after incubation with 20 µM EdU, as described in the methods. (A) Representative flow cytograms of each isolate grown either with or without EdU. Isolates grown without EdU were used to setup our gating strategy. The horizontal line marks the top of the fluorescent population in the sample grown without EdU. The vertical line separates bacterial cells on the right and debris on the left. As such, everything below the horizontal line and to the right (Quadrant 3; Q3) represents the unlabeled, or EdU-negative (EdU-), population, whereas anything above it and to the right (Quadrant 2; Q2) represents the EdU+ population (B) Growth curves of all isolates shown in (A) during incubation with EdU. (C-D). Growth rates for each of the (C) nine or (D) eight (excluding *B. infantis*) bacterial isolates were calculated as the steepest slope of their exponential growth curve (Max V) and were correlated with percent of cells labeled with EdU (Pearson correlation). *B. longum* = *Bifidobacterium longum* subspecies *infantis; B. caccae* = *Bacteroides caccae; B. fragilis* = *Bacteroides fragilis; P. intermedia* = *Prevotella intermedia; C. beijerinckii* = *Clostridium beijerinckii; E. faecalis* = *Enterococcus faecalis; R. bromii* = *Ruminococcus bromii; E. coli* = *Escherichia coli; A. muciniphila* = *Akkermansia muciniphila*.

**Figure S2.**
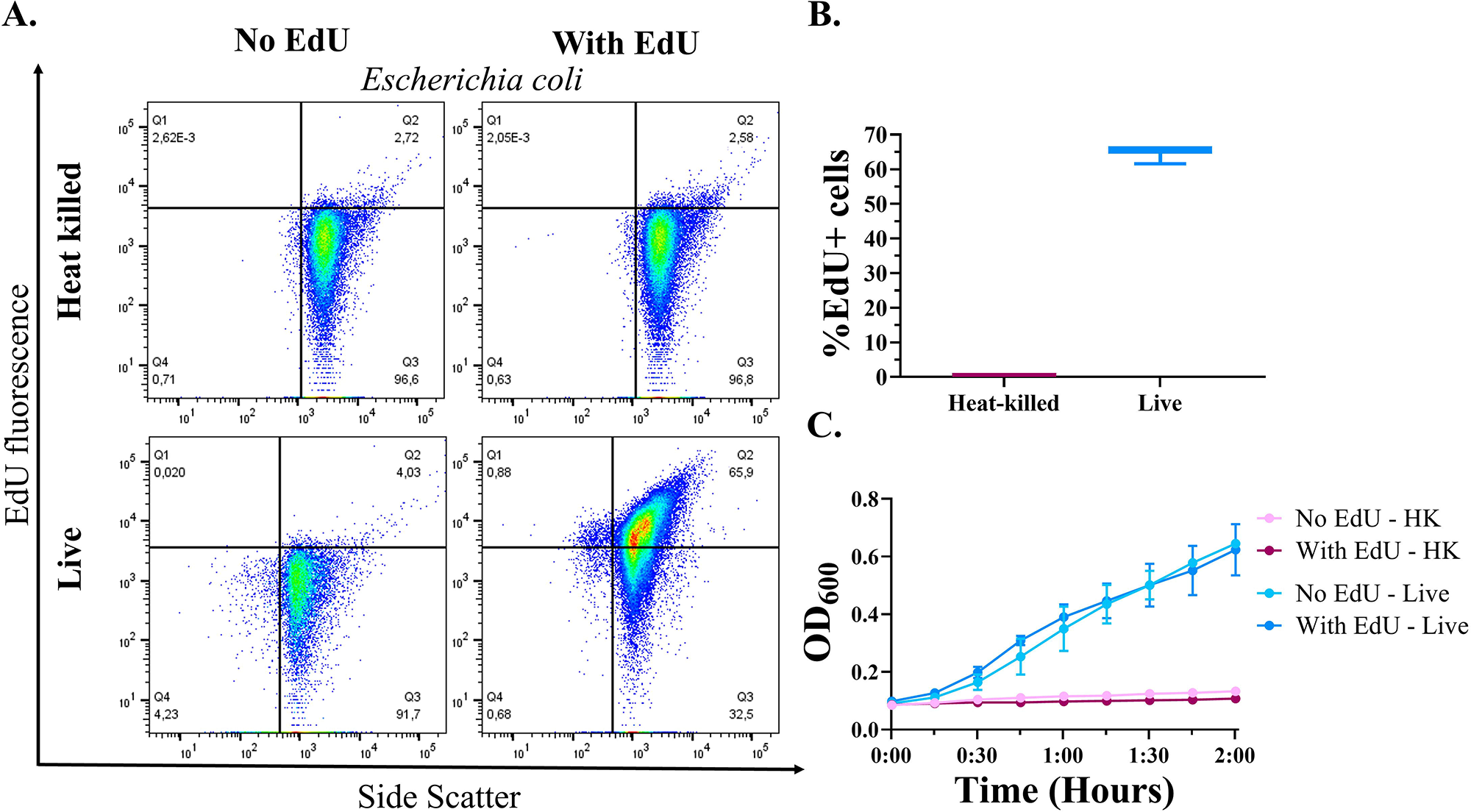
*Heat-killed* Escherichia coli *cells do not label with EdU*. *Escherichia coli* K12 cultures which were either heat-killed (HK; 65°C for 1 hour) or live underwent anaerobic incubation with 0 or 100 uM EdU for 2 hours, followed by the click procedure as described in the main text. (A) Representative flow cytograms of Escherichia coli K12 cultures which were either heat-killed (Top panel) or not (Bottom panel) before undergoing EdU-click. (B) Quantified proportion of EdU+ cells, heat-killed (HK) or live, after undergoing EdU-click (C) Growth curves of all conditions in (A).

**Figure S3.**
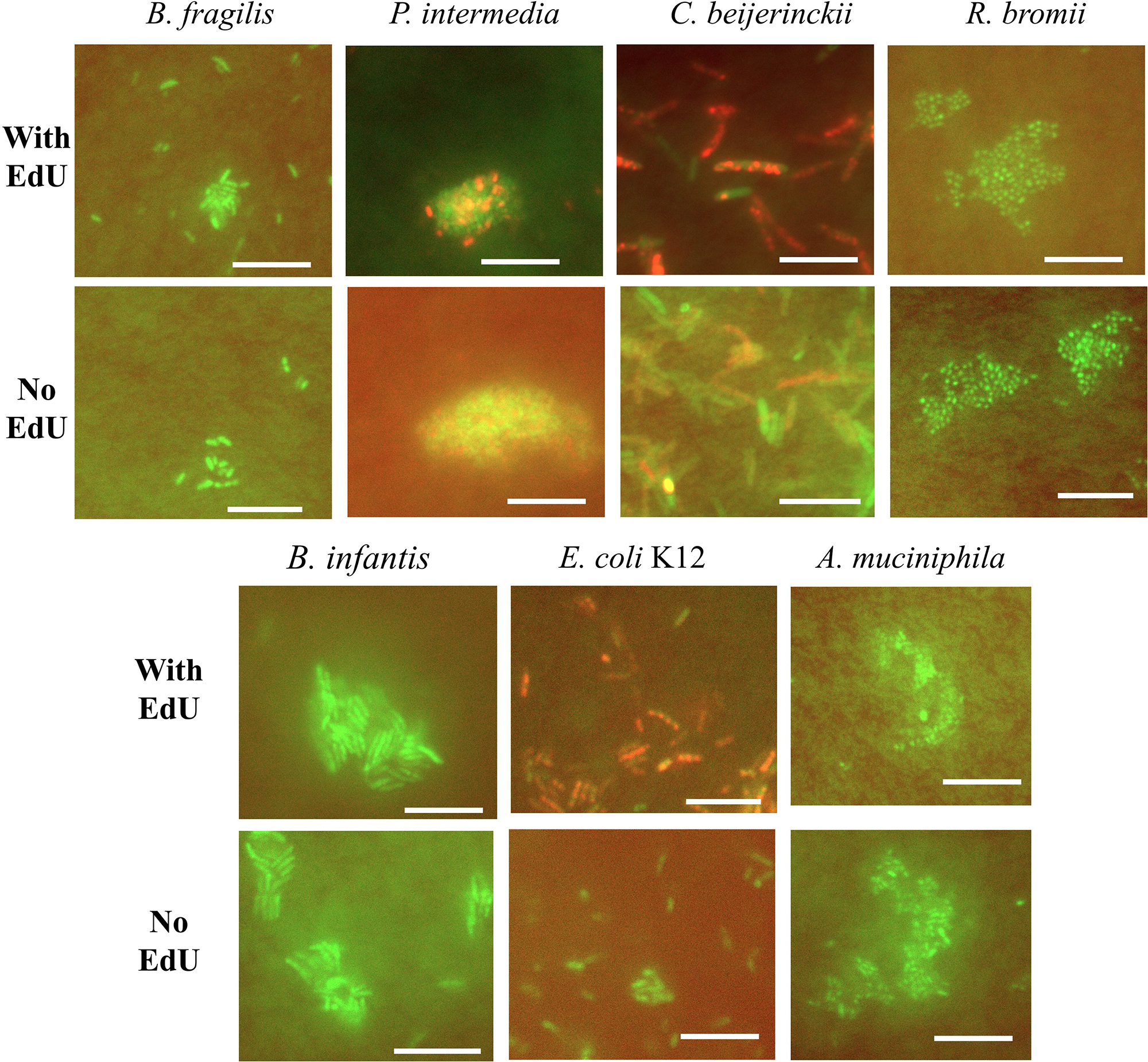
Epifluorescence microscopy of gut bacterial isolates after EdU-click. Human gut bacterial isolates which underwent EdU-click with or without EdU were stained with Sybr Green I nucleic acid dye and visualized using epifluorescence microscopy. Green = Sybr Green I, staining all bacterial nucleic acid content; Red = Alexa Fluor 647 azide, staining only newly replicated bacterial DNA. Any red signal seen in samples incubated without EdU was considered background noise, since these signals did not correspond to the location of bacterial cells as determined by Sybr Green I staining. Representative images of each isolate are shown. *B. fragilis = Bacteroides fragilis; P. intermedia = Prevotella intermedia; C. beijerinckii = Clostridium beijerinckii; R. bromii = Ruminococcus bromii; B. infantis = Bifidobacterium longum* subspecies *infantis; E. coli* K12 *= Escherichia coli* K12*; A. muciniphila = Akkermansia muciniphila*. Scale bar shows a width of 10 microns.

**Figure S4.**
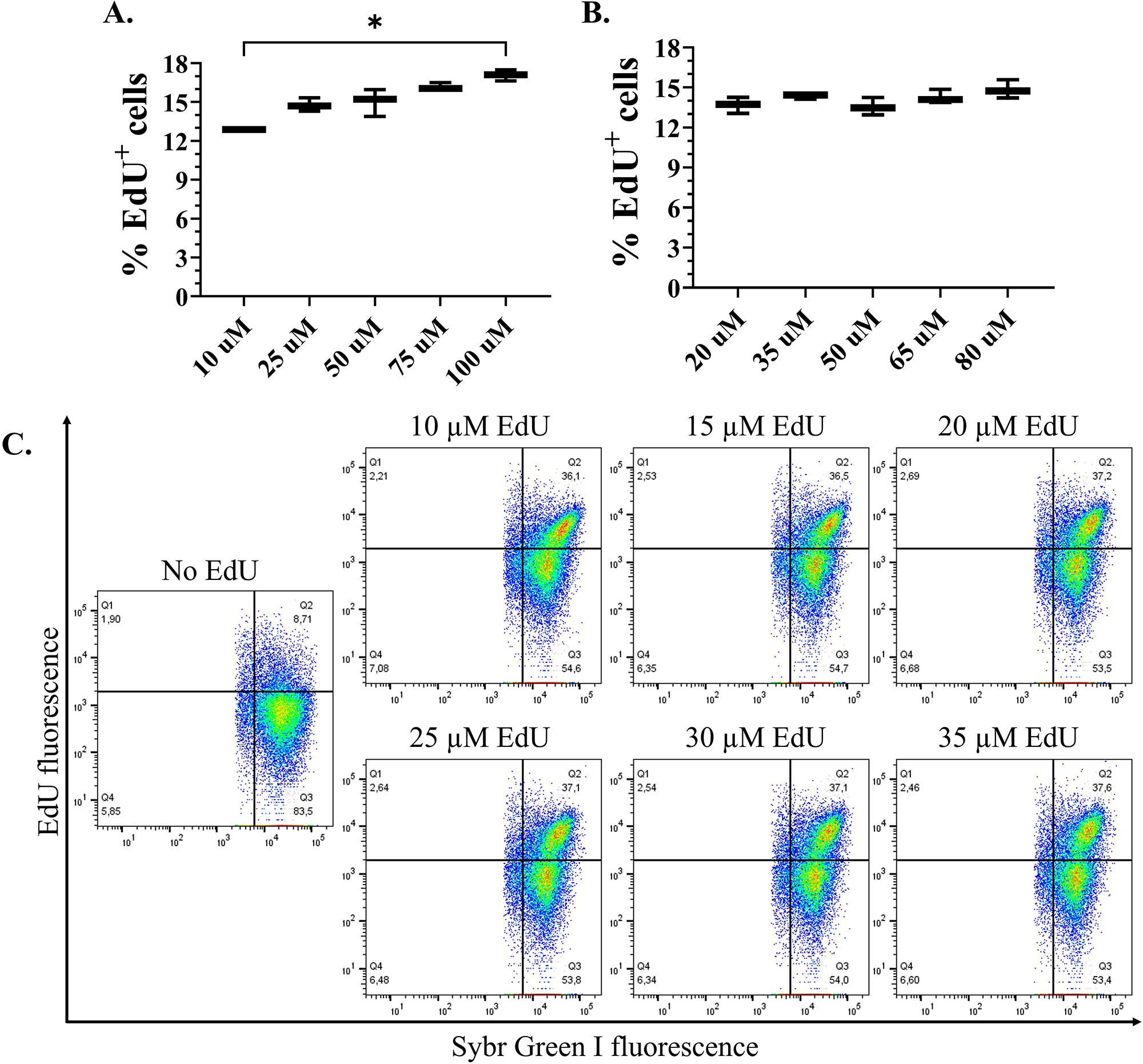
Optimization of EdU-click: EdU concentration. Mouse fecal bacteria from two independent experiments underwent EdU-click with varying concentrations of EdU and proportions of labeled cells were quantified using flow cytometry. (A) Proportions of EdU+ cells for EdU concentrations from 10 to 100 µM in 15 or 25 µM increments (B) Proportions of EdU+ cells for EdU concentrations from 20 to 80 µM in 15 µM increments. Statistical significance determined by Kruskal-Wallis with Dunn’s test to correct for multiple comparisons. Comparisons are considered significant at p ≤ 0.05. (C) Flow cytograms for fecal mouse bacteria incubated with concentrations of EdU from 10-35 µM.

**Figure S5.**
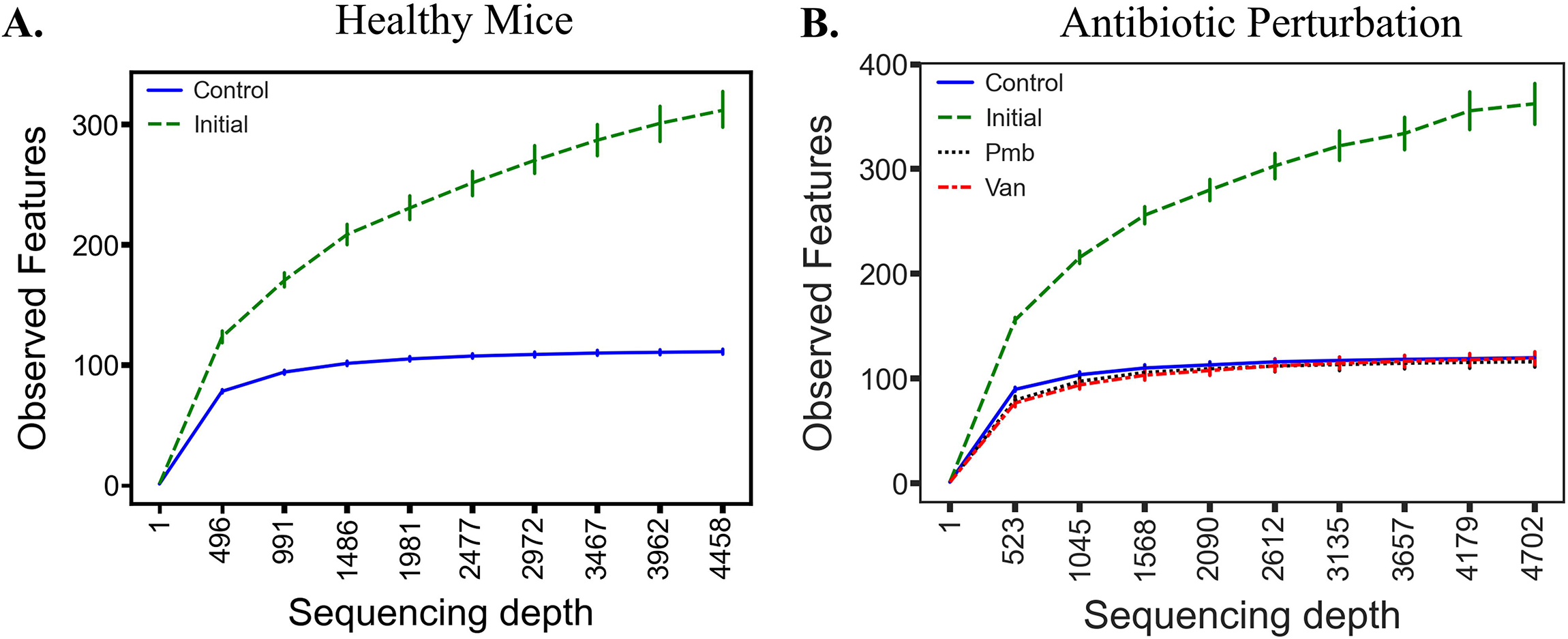
Rarefaction curves on the number of observed features in each experiment. The plots below display the rarefaction curves on the number of observed features for (A) the experiment on healthy mice (“Healthy Mice”) and (B) the antibiotic perturbation experiment (“Antibiotic Perturbation”), per treatment: Control, Initial; Van = vancomycin, Pmb = polymyxin B. The maximum sequencing depth in each plot was set at the depth at which these sequences were rarefied for alpha and beta diversity analyses.

**Figure S6.**
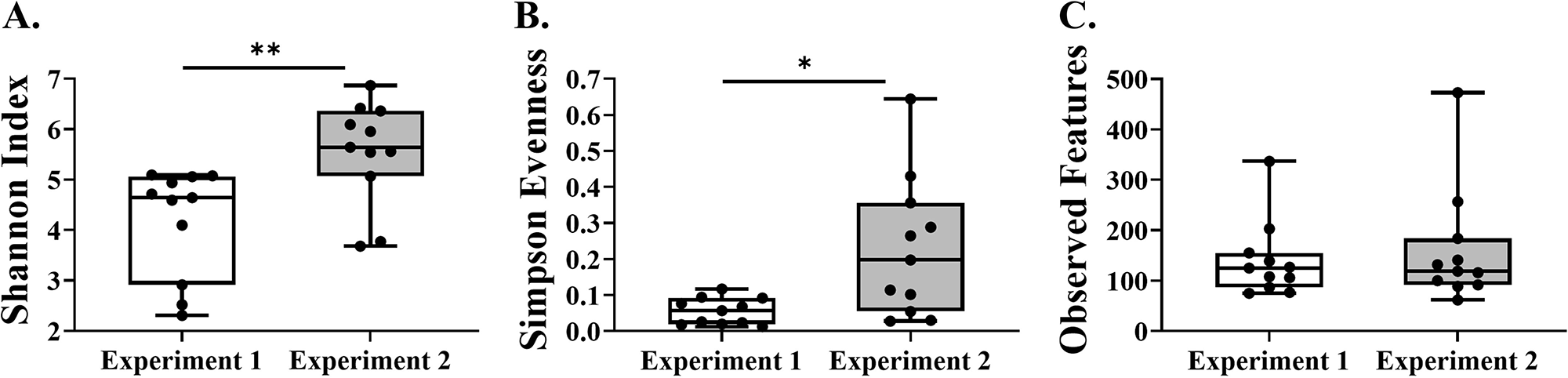
Alpha diversity metrics of healthy mice in two independent experiments. Within-sample alpha diversity plots of the bacterial communities from healthy C57BL/6 mice are shown below, calculated using Shannon’s index (A), Simpson Evenness (B), and Observed Features (C) for each experiment. The results from all the sorted fractions (initial and whole community, EdU+ and EdU-cells) are combined in these metrics.

**Table SI.** Characteristics of mouse fecal-derived bacterial growth curves when exposed to antibiotics. Mouse fecal-derived bacteria were incubated with or without a cell-membrane specific antibiotic, vancomycin or polymyxin B. The growth curves can be seen in Figure 4a. (A) Numerical summaries of some key characteristics of those growth curves are summarized here. Lag time is here defined as the time it takes until the bacterial community begins its exponential growth phase. Max V = maximum velocity, or maximum replication rate; AUC = area under the curve. (B) P-values comparing Condition 1 versus Condition 2 for all growth curve characteristics. All values were calculated in the EPOCH2 software and statistical analyses were done in GraphPad Prism using a one-way ANOVA with Dunnett’s test to correct for multiple comparisons, comparing the control to each treatment.

| Condition | Replicate | Lag Time<br>(HH:MM:SS) | Max V | AUC |
| --- | --- | --- | --- | --- |
| Control | 1 | 00:23:43 | 8.375 | 2.844 |
|  | 2 | 00:31:19 | 8.143 | 2.478 |
|  | 3 | 00:27:58 | 9.039 | 2.701 |
| Vancomycin | 1 | 00:45:07 | 7.607 | 2.223 |
|  | 2 | 00:40:07 | 8.935 | 2.359 |
|  | 3 | 00:35:07 | 7.214 | 2.346 |
| Polymyxin B | 1 | 01:25:07 | 3.011 | 1.683 |
|  | 2 | 00:55:07 | 6.587 | 1.958 |
|  | 3 | 01:00:07 | 5.785 | 2.002 |

Table SIB
| Condition 1 | Condition 2 | Lag Time | Max V | AUC |
| --- | --- | --- | --- | --- |
| Control | Vancomycin | P = 0.8807 | P = 0.7836 | **P = 0.0455** |
|  | Polymyxin B | **P = 0.0073** | **P = 0.0264** | **P = 0.0013** |

**Table SII.** Potential vancomycin resistance genes in an Ileibacterium valens *strain*. The protein sequences of a publicly available genome for *Ileibacterium valens* (see the main text) were run through the Comprehensive Antibiotic Resistance Database (CARD) Resistance Gene Identifier (RGI) to detect antibiotic resistance genes. Only those for vancomycin and related glycopeptide antibiotics are shown. Note that all hits shown are classified as “Loose” hits.

| **Antibiotic**<br>**Resistance**<br>**Ontology**<br>**(ARO)**<br>**Term** | **Antimicrobial**<br>**Resistance**<br>**(AMR) Gene**<br>**Family** | **Drug Class** | **Resistance**<br>**Mechanism** | **%**<br>**Identity**<br>**of**<br>**Matching**<br>**Region** | **%**<br>**Length of**<br>**Reference**<br>**Sequence** |
| --- | --- | --- | --- | --- | --- |
| vanN | glycopeptide<br>resistance gene<br>cluster, Van ligase | glycopeptide<br>antibiotic | antibiotic target<br>alteration | 29.41 | 121.28 |
| vanG | glycopeptide<br>resistance gene<br>cluster, Van ligase | glycopeptide<br>antibiotic | antibiotic target<br>alteration | 40.34 | 100.57 |
| vanK gene in<br>vanI cluster | glycopeptide<br>resistance gene<br>cluster, vanK | glycopeptide<br>antibiotic | antibiotic target<br>alteration | 25.29 | 103.73 |
| vanR gene in<br>vanA cluster | glycopeptide<br>resistance gene<br>cluster, vanR | glycopeptide<br>antibiotic | antibiotic target<br>alteration | 29.9 | 45.02 |
| vanR gene in<br>vanF cluster | glycopeptide<br>resistance gene<br>cluster, vanR | glycopeptide<br>antibiotic | antibiotic target<br>alteration | 41.67 | 102.60 |
| vanR gene in<br>vanB cluster | glycopeptide<br>resistance gene<br>cluster, vanR | glycopeptide<br>antibiotic | antibiotic target<br>alteration | 29.79 | 75.45 |
| vanR gene in<br>vanF cluster | glycopeptide<br>resistance gene<br>cluster, vanR | glycopeptide<br>antibiotic | antibiotic target<br>alteration | 30.7 | 97.40 |
| vanR gene in<br>vanL cluster | glycopeptide<br>resistance gene<br>cluster, vanR | glycopeptide<br>antibiotic | antibiotic target<br>alteration | 34.82 | 94.81 |
| vanR gene in<br>vanF cluster | glycopeptide<br>resistance gene<br>cluster, vanR | glycopeptide<br>antibiotic | antibiotic target<br>alteration | 40.0 | 103.90 |
| vanR gene in<br>vanN cluster | glycopeptide<br>resistance gene<br>cluster, vanR | glycopeptide<br>antibiotic | antibiotic target<br>alteration | 34.65 | 96.52 |
| vanT gene in<br>vanG cluster | glycopeptide<br>resistance gene<br>cluster, vanT | glycopeptide<br>antibiotic | antibiotic target<br>alteration | 38.27 | 55.06 |
| vanT gene in<br>vanG cluster | glycopeptide<br>resistance gene<br>cluster, vanT | glycopeptide<br>antibiotic | antibiotic target<br>alteration | 36.12 | 47.47 |
| vanU gene in<br>vanG cluster | glycopeptide<br>resistance gene<br>cluster, vanU | glycopeptide<br>antibiotic | antibiotic target<br>alteration | 35.48 | 97.33 |
| vanH gene in<br>vanB cluster | vanH,<br>glycopeptide<br>resistance gene<br>cluster | glycopeptide<br>antibiotic | antibiotic target<br>alteration | 26.14 | 119.81 |
| vanH gene in<br>vanA cluster | vanH,<br>glycopeptide<br>resistance gene<br>cluster | glycopeptide<br>antibiotic | antibiotic target<br>alteration | 35.29 | 88.20 |
| vanS gene in<br>vanF cluster | vanS,<br>glycopeptide<br>resistance gene<br>cluster | glycopeptide<br>antibiotic | antibiotic target<br>alteration | 28.14 | 208.77 |
| vanS gene in<br>vanM cluster | vanS,<br>glycopeptide<br>resistance gene<br>cluster | glycopeptide<br>antibiotic | antibiotic target<br>alteration | 27.95 | 196.76 |
| vanS gene in<br>vanF cluster | vanS,<br>glycopeptide resistance gene<br>cluster | glycopeptide<br>antibiotic | antibiotic target<br>alteration | 22.01 | 89.04 |
| vanY gene in<br>vanF cluster | vanY,<br>glycopeptide<br>resistance gene<br>cluster | glycopeptide<br>antibiotic | antibiotic target<br>alteration | 29.09 | 114.33 |
| vanY gene in<br>vanF cluster | vanY,<br>glycopeptide<br>resistance gene<br>cluster | glycopeptide<br>antibiotic | antibiotic target<br>alteration | 41.48 | 110.92 |
| vanZ gene in<br>vanF cluster | vanZ,<br>glycopeptide<br>resistance gene<br>cluster | glycopeptide<br>antibiotic | antibiotic target<br>alteration | 28.49 | 178.64 |

## References

1. Li J, Butcher J, Mack D, et al. Functional impacts of the intestinal microbiome in the pathogenesis of inflammatory bowel disease. Inflamm Bowel Dis. 2015 Jan;21(1):139–53.

2. Lloyd-Price J, Abu-Ali G, Huttenhower C. The healthy human microbiome. Genome Med. 2016 Apr 27;8(1):51.

3. Mirzaei MK, Maurice CF. Menage a trois in the human gut: interactions between host, bacteria and phages. Nat Rev Microbiol. 2017 Jul;15(7):397–408.

4. Shkoporov AN, Hill C. Bacteriophages of the Human Gut: The “Known Unknown” of the Microbiome. Cell Host Microbe. 2019 Feb 13;25(2):195–209.

5. Gill SR, Pop M, Deboy RT, et al. Metagenomic analysis of the human distal gut microbiome. Science. 2006 Jun 2;312(5778):1355–9.

6. Lozupone CA, Stombaugh JI, Gordon JI, et al. Diversity, stability and resilience of the human gut microbiota. Nature. 2012 Sep 13;489(7415):220–30.

7. Schirmer M, Garner A, Vlamakis H, et al. Microbial genes and pathways in inflammatory bowel disease. Nat Rev Microbiol. 2019 Aug;17(8):497–511.

8. Franzosa EA, Morgan XC, Segata N, et al. Relating the metatranscriptome and metagenome of the human gut. Proc Natl Acad Sci U S A. 2014 Jun 3;111(22):E2329–38.

9. Franzosa EA, Sirota-Madi A, Avila-Pacheco J, et al. Gut microbiome structure and metabolic activity in inflammatory bowel disease. Nat Microbiol. 2019 Feb;4(2):293–305.

10. Qin J, Li R, Raes J, et al. A human gut microbial gene catalogue established by metagenomic sequencing. Nature. 2010 Mar 4;464(7285):59–65.

11. Schirmer M, Franzosa EA, Lloyd-Price J, et al. Dynamics of metatranscription in the inflammatory bowel disease gut microbiome. Nat Microbiol. 2018 Mar;3(3):337–346.

12. Tropini C, Earle KA, Huang KC, et al. The Gut Microbiome: Connecting Spatial Organization to Function. Cell Host Microbe. 2017 Apr 12;21(4):433–442.

13. Moen AEF, Lindstrom JC, Tannaes TM, et al. The prevalence and transcriptional activity of the mucosal microbiota of ulcerative colitis patients. Sci Rep. 2018 Nov 22;8(1):17278.

14. Hatzenpichler R, Krukenberg V, Spietz RL, et al. Next-generation physiology approaches to study microbiome function at single cell level. Nat Rev Microbiol. 2020 Apr;18(4):241–256.

15. Brown CT, Olm MR, Thomas BC, et al. Measurement of bacterial replication rates in microbial communities. Nat Biotechnol. 2016 Dec;34(12):1256–1263.

16. Koch BJ, McHugh TA, Hayer M, et al. Estimating taxon-specific population dynamics in diverse microbial communities. Ecosphere. 2018;9(1):e02090.

17. Long AM, Hou S, Ignacio-Espinoza JC, et al. Benchmarking microbial growth rate predictions from metagenomes. ISME J. 2021 Jan;15(1):183–195.

18. Myhrvold C, Kotula JW, Hicks WM, et al. A distributed cell division counter reveals growth dynamics in the gut microbiota. Nat Commun. 2015 Nov 30;6:10039.

19. Riglar DT, Richmond DL, Potvin-Trottier L, et al. Bacterial variability in the mammalian gut captured by a single-cell synthetic oscillator. Nat Commun. 2019 Oct 11;10(1):4665.

20. Joseph TA, Chlenski P, Litman A, et al. Accurate and robust inference of microbial growth dynamics from metagenomic sequencing reveals personalized growth rates. Genome Res. 2022 Mar;32(3):558–568.

21. Korem T, Zeevi D, Suez J, et al. Growth dynamics of gut microbiota in health and disease inferred from single metagenomic samples. Science. 2015 Sep 4;349(6252):1101–1106.

22. Hudak JE, Alvarez D, Skelly A, et al. Illuminating vital surface molecules of symbionts in health and disease. Nat Microbiol. 2017 Jun 26;2:17099.

23. Kuru E, Hughes HV, Brown PJ, et al. In Situ probing of newly synthesized peptidoglycan in live bacteria with fluorescent D-amino acids. Angew Chem Int Ed Engl. 2012 Dec 7;51(50):12519–23.

24. Lin L, Wu Q, Song J, et al. Revealing the in vivo growth and division patterns of mouse gut bacteria. Sci Adv. 2020 Sep;6(36).

25. Wang W, Lin L, Du Y, et al. Assessing the viability of transplanted gut microbiota by sequential tagging with D-amino acid-based metabolic probes. Nat Commun. 2019 Mar 21;10(1):1317.

26. Taguer M, Shapiro BJ, Maurice CF. Translational activity is uncoupled from nucleic acid content in bacterial cells of the human gut microbiota. Gut Microbes. 2021 Jan-Dec;13(1):1ߝ15.

27. Salic A, Mitchison TJ. A chemical method for fast and sensitive detection of DNA synthesis in vivo. Proc Natl Acad Sci U S A. 2008 Feb 19;105(7):2415–20.

28. Steward GF, Azam F. Bromodeoxyuridine as an alternative to 3H-thymidine for measuring bacterial productivity in aquatic samples. Aquatic Microbial Ecology. 1999;19:57–66.

29. Smriga S, Samo T, Malfatti F, et al. Individual cell DNA synthesis within natural marine bacterial assemblages as detected by ‘click’ chemistry. Aquatic Microbial Ecology. 2014 07/10;72:269–280.

30. Kurm V, van der Putten WH, de Boer W, et al. Low abundant soil bacteria can be metabolically versatile and fast growing. Ecology. 2017;98(2):555–564.

31. Ferullo DJ, Cooper DL, Moore HR, et al. Cell cycle synchronization of Escherichia coli using the stringent response, with fluorescence labeling assays for DNA content and replication. Methods. 2009 May;48(1):8–13.

32. Spahn C, Endesfelder U, Heilemann M. Super-resolution imaging of Escherichia coli nucleoids reveals highly structured and asymmetric segregation during fast growth. J Struct Biol. 2014 Mar;185(3):243–9.

33. Rousk J, Baath E. Growth of saprotrophic fungi and bacteria in soil. FEMS Microbiol Ecol. 2011 Oct;78(1):17–30.

34. Singh G, Brass A, Cruickshank SM, et al. Cage and maternal effects on the bacterial communities of the murine gut. Sci Rep. 2021 May 10;11(1):9841.

35. Jeffrey WH, Paul JH. Thymidine uptake, thymidine incorporation, and thymidine kinase activity in marine bacterium isolates. Appl Environ Microbiol. 1990;56(5):1367–1372.

36. Pérez MT, Hörtnagl P, Sommaruga R. Contrasting ability to take up leucine and thymidine among freshwater bacterial groups: implications for bacterial production measurements. Environmental Microbiology. 2010;12(1):74–82.

37. Pedrós-Alió C, Newell SY. Microautoradiography study of thymidine uptake in brackish waters around Sapelo Island, Georgia, USA. Marine Ecology-progress Series – MAR ECOL-PROGR SER. 1989 07/06;55:83–94.

38. Tsuchiya K, Sano T, Tomioka N, et al. Incorporation characteristics of exogenous 15N-labeled thymidine, deoxyadenosine, deoxyguanosine and deoxycytidine into bacterial DNA. PLOS ONE. 2020;15(2):e0229740.

39. Cox LM, Sohn J, Tyrrell KL, et al. Description of two novel members of the family Erysipelotrichaceae: Ileibacterium valens gen. nov., sp. nov. and Dubosiella newyorkensis, gen. nov., sp. nov., from the murine intestine, and emendation to the description of Faecalibaculum rodentium. Int J Syst Evol Microbiol. 2017 May;67(5):1247–1254.

40. Bhat SF, Pinney SE, Kennedy KM, et al. Exposure to high fructose corn syrup during adolescence in the mouse alters hepatic metabolism and the microbiome in a sex-specific manner. J Physiol. 2021 Mar;599(5):1487–1511.

41. den Hartigh LJ, Gao Z, Goodspeed L, et al. Obese Mice Losing Weight Due to trans-10,cis-12 Conjugated Linoleic Acid Supplementation or Food Restriction Harbor Distinct Gut Microbiota. J Nutr. 2018 Apr 1;148(4):562–572.

42. Xia F, Xiang S, Chen Z, et al. The probiotic effects of AB23A on high-fat-diet-induced non-alcoholic fatty liver disease in mice may be associated with suppressing the serum levels of lipopolysaccharides and branched-chain amino acids. Arch Biochem Biophys. 2021 Dec 15;714:109080.

43. Weissman JL, Hou S, Fuhrman JA. Estimating maximal microbial growth rates from cultures, metagenomes, and single cells via codon usage patterns. Proc Natl Acad Sci U S A. 2021 Mar 23;118(12).

44. Alcock BP, Raphenya AR, Lau TTY, et al. CARD 2020: antibiotic resistome surveillance with the comprehensive antibiotic resistance database. Nucleic Acids Res. 2020 Jan 8;48(D1):D517–D525.

45. Kwok AHY, Li Y, Jiang J, et al. Complete genome assembly and characterization of an outbreak strain of the causative agent of swine erysipelas--Erysipelothrix rhusiopathiae SY1027. BMC Microbiol. 2014;14:176–176.

46. Romney M, Cheung S, Montessori V. Erysipelothrix rhusiopathiae endocarditis and presumed osteomyelitis. Can J Infect Dis. 2001 Jul;12(4):254–6.

47. Wu J, Liu M, Zhou M, et al. Isolation and genomic characterization of five novel strains of Erysipelotrichaceae from commercial pigs. BMC Microbiol. 2021 Apr 23;21(1):125.

48. Klein G, Hallmann C, Casas IA, et al. Exclusion of vanA, vanB and vanC type glycopeptide resistance in strains of Lactobacillus reuteri and Lactobacillus rhamnosus used as probiotics by polymerase chain reaction and hybridization methods. J Appl Microbiol. 2000 Nov;89(5):815–24.

49. Bolyen E, Rideout JR, Dillon MR, et al. Reproducible, interactive, scalable and extensible microbiome data science using QIIME 2. Nat Biotechnol. 2019 Aug;37(8):852–857.

50. Kaul A, Mandal S, Davidov O, et al. Analysis of Microbiome Data in the Presence of Excess Zeros. Front Microbiol. 2017;8:2114.

